# Urbanization and wealth alter arthropod biodiversity and pollination services in community gardens

**DOI:** 10.64898/2026.09.18.752681

**Authors:** Asia Kaiser, Noah Mayer, Rene Aronson, Julian Resasco

**Affiliations:** Department of Ecology and Evolutionary Biology, The University of Colorado Boulder, Boulder, CO, USA

**Keywords:** urban biodiversity, socioecological, pollinators, insects, urban agriculture, cucumbers

## Abstract

1. As city populations expand globally, understanding how urbanization shapes ecosystem services is increasingly important. Urban agriculture plays a key role in food access, particularly in low-income areas with limited access to fresh produce. Arthropods play important roles in these systems through pollination, herbivory, pest control, and nutrient cycling. Urban development of natural areas often reduces biodiversity, including that of arthropods. Additionally, urban biodiversity frequently correlates positively with socioeconomic conditions, a pattern known as the luxury effect.
2. In this study, we examined how gradients in urbanization and socioeconomic context relate to local and landscape habitat quality, arthropod abundance, and pollination services across 21 community gardens in the Denver, CO Metro Area. To quantify ecosystem services, we employed a sentinel plant, *Cucumis sativus* (cucumber), and collected arthropods (insects, myriapods, arachnids, isopods), assigning them to functional groups based on their known feeding behavior. We evaluated direct and indirect pathways linking urban matrix characteristics to arthropod-mediated ecosystem services.
3. The abundance of all arthropod groups (including pollinators, herbivores, and natural enemies) declined steeply with increasing impervious surface. Neighborhood wealth was associated only with bee species richness and Shannon diversity and displayed a hump-shaped pattern that peaked at wealth levels 35-46% above the national average, then declined. Multivariate GLMs revealed that Western honeybees (*Apis mellifera*) were the only species driving differences in bee community composition, with their abundance positively related to neighborhood wealth.
4. We also observed pollen limitation in our cucumber plants, and as bee Shannon diversity increased, cucumber production increased. Environmental covariates alone showed no direct relationship with cucumber production, indicating that insect pollination mediated this effect. This work identifies the biophysical and social-ecological drivers of urban arthropod biodiversity and ecosystem services, revealing where crop production may be most vulnerable to decline in cities.

## INTRODUCTION

Urban community gardens provide a variety of benefits from nature to city dwellers. They offer material and provisioning services through produce and flowers, as well as non-material cultural services through recreation, community engagement, and access to nature (Díaz et al., 2018; Drake & Lawson, 2015). Previous studies have found that most plants grown in gardens are edible crops, with garden outputs providing a substantial portion of household produce needs. Low-income households often acquire a higher percentage of their household produce through these garden crops and prioritize growing edible over ornamental crop species (Clarke & Jenerette, 2015; Gregory et al., 2016). A national survey of over 8,000 community garden users across all fifty U.S. states found that more than 99% of respondents reported “food production and access” and “nutrition/improved diet” as benefits of their community garden, with 96% also citing “environmental benefits” (Drake & Lawson, 2015). Urban community gardens, therefore, play a critical role in improving food security and access to nutritious food, particularly in low-income areas with limited access to fresh produce (Krishnan et al., 2016).

Urban agroecosystems rely on services provided by arthropods, including pollination, natural enemy pest control, and nutrient cycling, to support ecosystem functioning. A meta-analysis of urban agriculture found that 27% of crop species in urban agroecosystems, including melons, pumpkins, peppers, cucumbers, and tomatoes, depend on insect pollination (Nascimento et al., 2025). Herbivore damage is also a major concern in urban agricultural systems, making natural enemy pest control equally important (Liere et al., 2020). A meta-analysis of 89 studies examining pollinators and natural enemies found that higher species richness and abundance enhance pollination and pest-control services (Dainese et al., 2019). Thus, arthropod abundance alone is not a sufficient indicator of ecosystem service value; taxonomic diversity must also be considered. Globally, population declines across multiple insect taxa have been reported (Wagner, 2020), and many pollinators face an elevated risk of extinction, with bees being the most at-risk pollinator group (Cornelisse et al., 2025). Despite growing evidence of global insect declines, substantial knowledge gaps remain regarding how urbanization specifically shapes arthropod diversity, community composition, and multitrophic interactions (Lewthwaite et al., 2024). Given their critical role in providing ecosystem services, understanding arthropod biodiversity in urban ecosystems is needed.

Urban community gardens not only benefit humans but can also serve as important refugia for arthropods by providing habitat and resources (Hall et al., 2017). A meta-analysis of biodiversity in community gardens found that gardens used primarily for food production support higher arthropod biodiversity than those dominated by ornamental plants, indicating that these systems can simultaneously benefit human livelihoods and arthropod conservation (Nascimento et al., 2025). Unfortunately, these conservation benefits to arthropods are not universal; while urban ecosystems may serve as a refuge for some taxa, they may be inhospitable for others (Baldock et al., 2019). This can lead to trait syndromes driven by environmental filtering of particular taxa (Hahs et al., 2023). Filtering of more sensitive species may alter arthropod community composition and influence ecological dynamics in community gardens, for example, by reducing predation services and increasing herbivory. Indeed, urban gardens have been shown to have lower densities of natural enemies and higher densities of herbivorous pests than natural areas outside cities, thereby hampering biological control in these systems (Gregory et al., 2016; Korányi et al., 2022). Higher agricultural land-use cover surrounding urban gardens has also been associated with lower natural enemy–to–herbivore ratios (Philpott et al., 2020). Landscape simplification has been shown to reduce pollination and pest-control services, primarily by decreasing arthropod species richness (Dainese et al., 2019; Kennedy et al., 2013). In addition, the benefits conferred to arthropods by increased green space in certain urban habitats may be offset by increased pesticide use, heavy irrigation, and changes in microhabitats (Guaitero et al., 2026; Knauer et al., 2025).

Socioeconomic variables have repeatedly been shown to correlate with biodiversity across taxa in urban environments (Chamberlain et al., 2019; Leong et al., 2018). Early studies reported a positive relationship between neighborhood wealth and biodiversity (Hope et al., 2003); yet more recent work suggests that this relationship may not hold across all biomes, across the full wealth spectrum, or specifically for arthropod taxa (Chamberlain et al., 2020; Kaiser & Resasco, 2024). Features common in wealthier neighborhoods—such as increased tree canopy cover, turfgrass, and ornamental plantings—may provide sufficient habitat and provisioning for larger animal taxa like birds and mammals but could negatively affect certain arthropod populations. For example, the richness and abundance of bees have been shown to be higher in sites with lower tree canopy cover (Hall et al., 2019). Furthermore, many arthropod species form specialist relationships with particular floral genera; if these host plants are outcompeted or intentionally excluded by homeowners in favor of exotic ornamentals, wealthier neighborhoods may host lower arthropod biodiversity (Kaiser & Resasco, 2024). It remains unknown what the underlying patterns are for arthropod biodiversity, or whether the socioeconomic factors shaping these communities also affect ecosystem services in urban agroecosystems, potentially altering crop yields. This has important implications for environmental justice with regard to equal access to nature and ecosystem services.

Here, we investigate how gradients in urbanization (measured by impervious surface area) and socioeconomic factors influence arthropod community composition (including pollinators, herbivores, and beneficial predators), and, consequently, the pollination and pest-control ecosystem services provided at community garden sites. We address two primary research questions: **(Q1)** How do arthropod diversity and community composition relate to biodiversity-mediated ecosystem services (pollination and pest control)? And **(Q2)** How do urban landscape and socioeconomic characteristics influence these patterns? We hypothesize that socioeconomic factors influence both local and landscape features in the matrix surrounding garden sites, thereby affecting ecosystem services and crop productivity by altering arthropod biodiversity and community composition. These local and landscape features will include floral resources, bare ground, tree cover, and impervious surface levels, which are predictors of matrix quality, at community garden sites and in the surrounding landscape. If this hypothesis is supported, then arthropod populations and, consequently, fruit set in pollinator-dependent plants should depend on habitat quality at garden sites and in the surrounding neighborhood. This work aims to enhance our understanding of urban agroecosystems and to identify areas at risk of insufficient ecosystem services and reduced crop yields.

## METHODS

### Study sites and socio-ecological landscape context

This study was conducted at 21 community gardens across five cities in the Denver Metro Area: Denver, Lakewood, Aurora, Golden, and Englewood, Colorado, USA (Table S1.1). The Denver Metro area is situated in the Front Range Urban Corridor, along the southeastern base of the Rocky Mountains, with a semi-arid climate. The 21 sites were all a part of the Denver Urban Gardens (DUG) Collective (https://dug.org/), which has over 190 different participating gardens. All sites use organic growing practices because DUG has a strict policy against chemical pesticides, herbicides, and chemical fertilizers. We selected a pool of sites for inclusion in the study by excluding private-access sites and sites with managed honeybee hives. To ensure that we had the full income spectrum represented in our site sample, we employed stratified random sampling and categorized sites into five strata by neighborhood wealth. We then randomly selected sites from each wealth stratum for inclusion in our study, ensuring they were at least 1.5 km apart (Fig. 1) and verified that wealth and impervious surface values (described below) were not correlated using a Pearson correlation test (t = −0.67, df = 19, p-value = 0.507).

**Figure 1.**
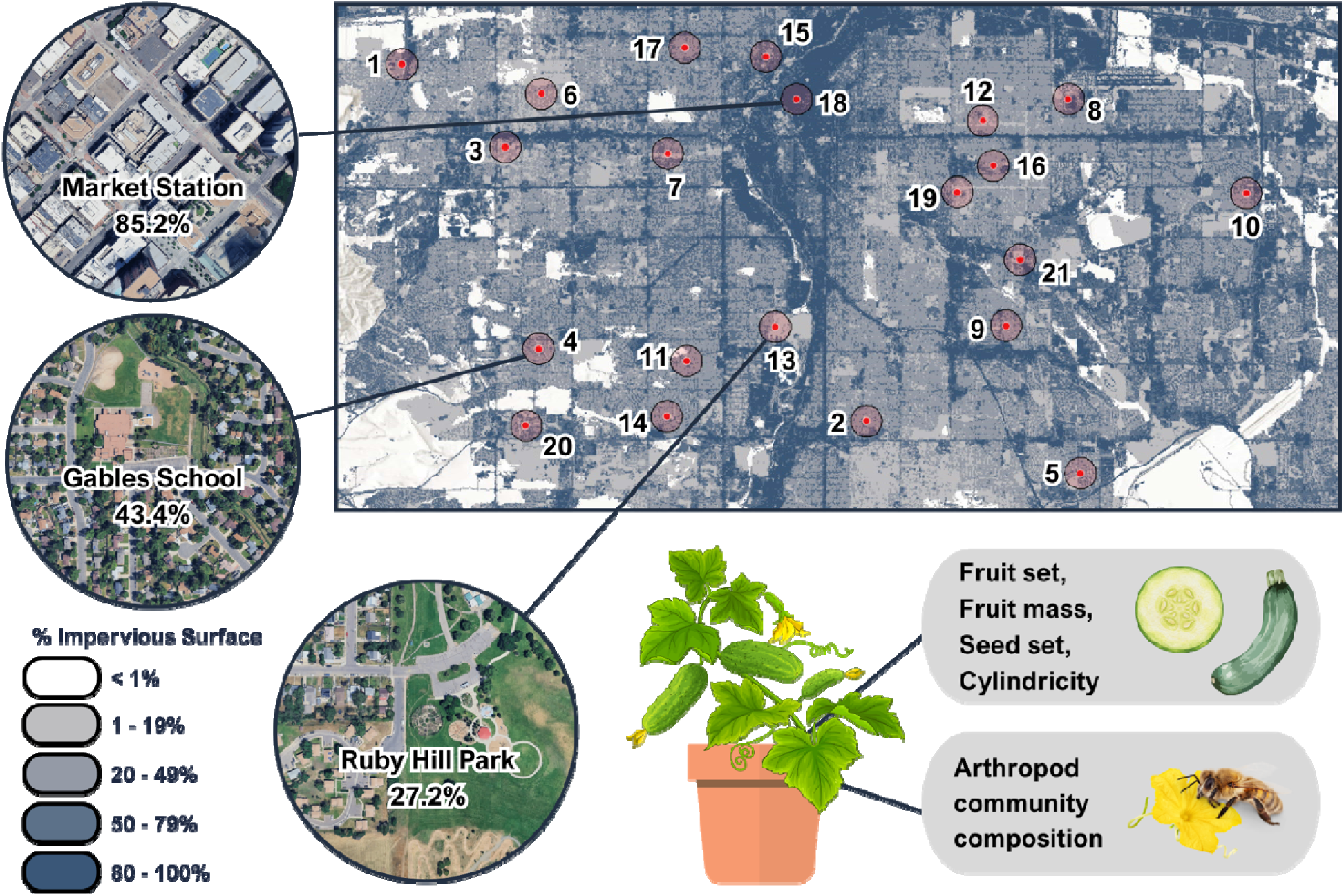
An imperviousness map of the Denver Metro Area showing 21 study sites and their 500-meter buffers. Numbers correspond with the garden names listed in Table S1.1. On the left, from top to bottom, are aerial photographs of sites with high, medium, and low imperviousness, and the percent imperviousness within a 200-meter buffer is listed. The bottom-right panel shows the response variables measured at each site: arthropod community composition and cucumber fruit mass, fruit set, and seed set. *Image Credits*: ESRI ArcGIS, Canva Pro License.

**Figure 2.**
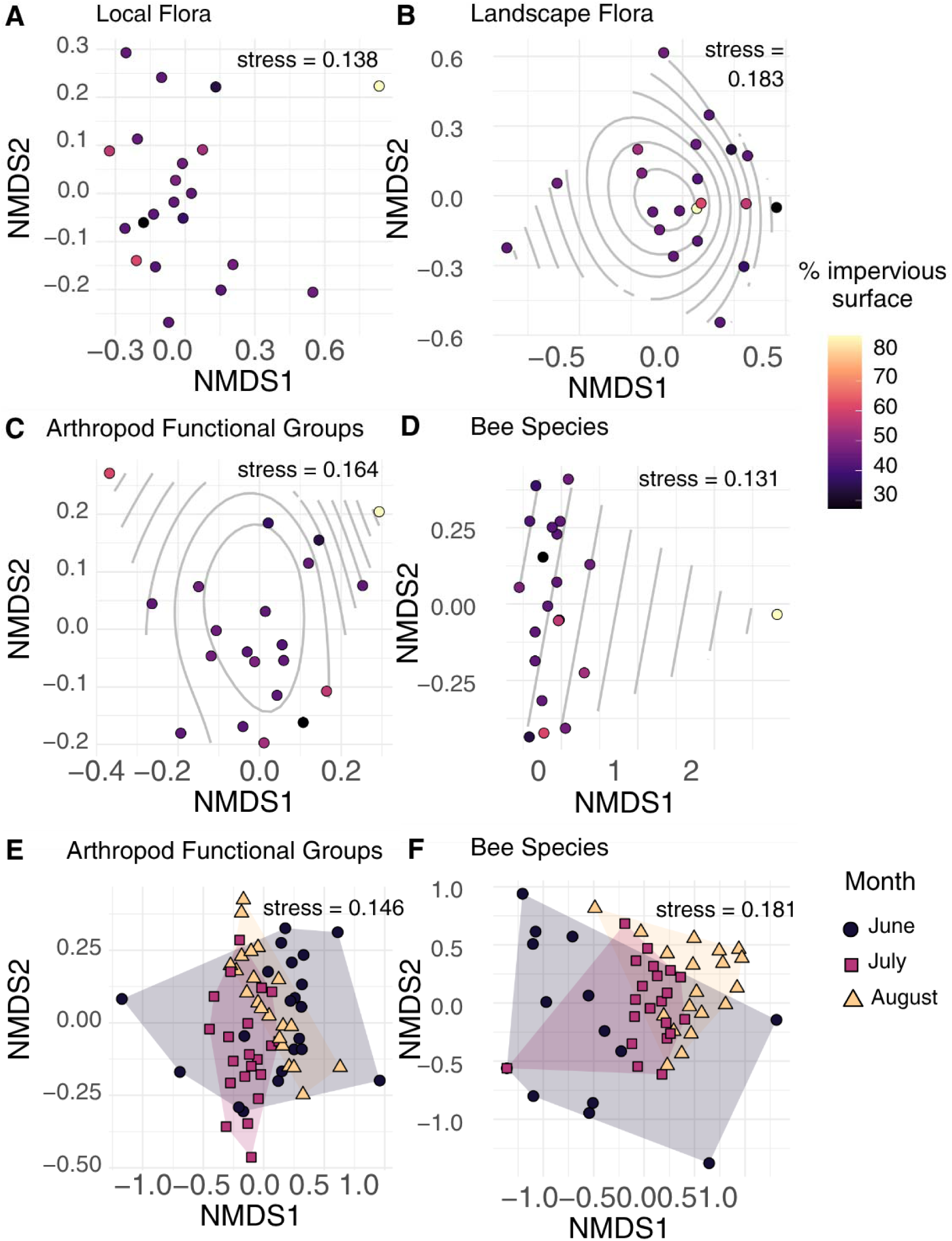
Non-metric multidimensional scaling (NMDS) ordinations of site-level community composition based on Bray–Curtis dissimilarities for **a)** local floral families observed within community gardens, **b)** landscape floral families recorded via iNaturalist within a 1 km buffer of each site, **c)** arthropod functional groups collected across the summer, and **d)** bee species collected across the summer. In panels **b–d**, contours represent smooth surfaces of impervious surface within 200 m, fitted using generalized additive models, where impervious surface was significantly associated with community composition. Panels **e)** and **f)** show NMDS ordinations of monthly arthropod functional group and bee species communities, respectively, with points grouped by month.

At the landscape scale, we calculated the mean degree of impervious surface and tree cover surrounding our sites from the Fractional Impervious Surface (2024) and Tree Canopy Cover (2023) 30-m^2^ resolution rasters from the National Land Cover Database (United States Geological Survey, 2025) with the R package sp (v2.2.1; Pebesma et al., 2026). The value for each pixel in these rasters represents the percentage of that 30 m^2^ covered by impervious surface (for the Fractional Impervious Surface raster) or tree canopy (for the Tree Canopy Cover raster). We calculated the mean raster values within 200-m and 500-m buffers around each site. At the 200-meter scale, fractional impervious surface values across our sites ranged from 27.20 to 85.16% (mean 46.31% ± SD 11.37%); tree canopy cover values ranged from 0.58 to 20.82% (mean 11.61% ± SD 5.13%; Table S1.1). To quantify the socioeconomic conditions surrounding our sites, we utilized ESRI’s 2025 Wealth Index, calculated within a 500-m buffer surrounding each site in ArcGIS. We selected a larger buffer (500 m) to account for the lower data density underlying the index at finer scales and the fact that Wealth Index is not consistently available for all sites at smaller spatial extents. This index encompasses household income, average net worth, and the value of possessions and resources to help identify urban areas with higher or lower standards of living. A value of 100 represents a wealth index comparable to the national average. The wealth index values across our sites ranged from 62 to 272 (mean 137 ± SD 58; Table S1.1). To assess additional landscape variables that may influence arthropod habitat quality, we extracted the percentages of water, wetland, shrub, and grassland within 1-km buffers around each site using the National Land Cover Database land-cover raster. Because water and wetland coverage, as well as shrub and grassland coverage, were minimal at this scale, we combined them into two broader classes: “water/wetland” and “shrub/grassland.” Although previous research suggests that smaller spatial scales are often more predictive for arthropods (Kaiser & Resasco, 2024), data for several cover types were too sparse at 200-m and 500-m scales, necessitating the broader 1-km buffer for our landscape analysis.

At the garden (local) scale, we collected data on floral abundance and diversity. To do this, we measured the total area in m^2^ of each garden site covered by flowering plants from July 30 to August 6, 2025. Flowering areas were delineated in the field using visual surveys and mapped onto site images. Floral area was then calculated in ImageJ. Within each garden plot at a site, we recorded the number of unique taxa. We identified them to the lowest possible taxonomic level in the field, using the iNaturalist taxa identifier tool and our own plant identification expertise. By creating a new observation for each taxon each time it appeared in a new garden plot, we were able to get a proxy measure of the local abundance of different plant taxa at each site. In addition, to obtain a proxy for floral richness in the landscape surrounding the gardens, we downloaded research-grade iNaturalist observations from the Global Biodiversity Information Facility (GBIF.org, 2026) within a 1-km buffer around each garden site. We selected this scale to ensure sufficient observations for calculating biodiversity statistics, to prevent site buffers from overlapping, and to match the scale of our other landscape predictors. To capture other local features shown to affect arthropod populations (Philpott et al., 2014), we also measured the amounts of wooded and shrubby areas, bare ground, patchy vegetation, and fine- and coarse- woody debris in m^2^ within each garden site, by measuring area using transect tape.

### Pollination Services

To measure pollination services, we used ‘Spacemaster cucumbers, a bush-type cultivar of *Cucumis sativus,* as a sentinel plant. We selected this cucumber variety to evaluate insect-mediated pollination services because it is monoecious, requiring cross-pollination between its male and female flowers (Jagger, 1937), and does not fruit in the absence of pollinating insects (Motzke et al., 2015). It is also commonly grown in community gardens, with 100% of our sites having gardeners growing cucumbers. We started the cucumbers from seed in a greenhouse on April 28, 2025, and then placed six plants at each site on June 11-12, 2025 (n = 126). At each site, three pairs of plants were placed in the general garden area. To test for pollen limitation, we manually pollinated one plant in each pair, leaving the other plant untreated to serve as an open-pollination comparison group. If open-pollinated plants produce less than hand-pollinated plants, this indicates reduced pollination services at the site. To hand-pollinate, we used a paintbrush and transferred pollen from male flowers to female flowers on the same plant. We visited each site twice a week to pollinate all open female flowers, water the base of the cucumber plants, and collect mature cucumbers. Garden leaders and volunteers assisted with regular watering of cucumber plants to prevent them from drying out. Cucumbers were collected when the base of the fruit began to change color from green to yellow but was still firm. Collected cucumbers were placed in a cooler and transported back to the lab, where we recorded their wet mass, length, width, volume, and seed count from a longitudinal cross-section. We also recorded a qualitative metric (yes/no) indicating whether the cucumber was cylindrical, a metric encompassing circularity and straightness, in accordance with the USDA cucumber grades and standards (USDA, 2016); example photographs in Fig. S1.1).

### Arthropod sampling

We sampled arthropods at each site using vane and pitfall traps in the field for 72 hours (± 2 hours) from June 17 to June 20, July 15 to July 18, August 11 to 14, and August 19 to 23, 2025. Collections for August were split into two time periods; in the first collection round, seven sites had vane traps that collapsed in the field and required resampling. We used 50 mL centrifuge tubes as pitfall traps and placed two in the field at least 10 meters apart during each collection event. One fluorescent blue vane trap was placed at each site, approximately 1.5 meters above ground, in direct sunlight. To preserve and capture specimens, both pitfall and vane traps were filled with a solution of distilled water and ethanol, with a drop of dish soap to reduce surface tension. After each collection round, we returned the traps to the lab to pin and identify specimens. All arthropods were identified to the lowest taxonomic level possible to assign a functional group and feeding habit (Table S1.2). Mites (Subclass: Acari) were not identified beyond order due to limited taxonomic expertise. Many flies (Order: Diptera, n = 198) were also left unidentified (with the exception of Syrphidae and Culicomorpha) because they rapidly deteriorated in our traps, which prevented reliable identification and functional group assignment. Most taxa were identified to order using a variety of identification guides (Discover Life, 2026; Dunford & Long, 2002; Gullan & Cranston, 2014) and local expert opinion from the Resasco Lab at the University of Colorado Boulder. Beetles and Hemiptera were identified to lower taxonomic levels (family or genus), as there is significant variability in dietary behavior within these clades, with some species serving as beneficial predators and others as crop-damaging pests. We also selected bees as a focal group to obtain species-level data on (Epifamily: Anthophila) because of their important roles as pollinators and their conservation value as a group (Cornelisse et al., 2025; Wenzel et al., 2020). Bees were identified to the species level using dichotomous keys and identification guides (Carril & Wilson, 2023; Danforth et al., 2024; Discover Life, 2026) and local expert opinion (Asia Kaiser, 2025; identifications in Table S1.3). All arthropods were assigned to a functional group (pollinator, herbivore, predator, omnivore, detritivore, parasitoid, granivore, hematophagous, scavenger, fungivore) based on their known feeding behavior and life stage at the time of capture (BugGuide, n.d.; Gullan & Cranston, 2014). Pollinators were classified based on floral visitation behavior rather than strictly on their trophic diet. Some specimens were too damaged/degraded to confidently identify to a lower taxonomic level (7.7% of data); therefore, their functional group was recorded as unknown (Table S1.2).

### Data Analyses

#### Local and Landscape Variables and Wealth Index

We tested the relationship between wealth index and each local habitat quality variable measured at garden sites (garden floral richness and Shannon diversity, shrub and tree cover, bare ground, and floral species per m²), as well as landscape variables within a 1-km buffer around sites (GBIF floral observation richness and Shannon diversity, water and wetland cover, shrub and grassland cover). Tree canopy cover within a 500-m radius was used as a landscape variable, as its values were higher than those of the other landscape variables, and this scale better aligned with the spatial resolution of the wealth index data. We evaluated floral diversity using two biodiversity metrics: species richness, which measures the total number of species, and the Shannon diversity index, which captures both species richness and evenness. GBIF floral observation richness and Shannon diversity were extrapolated using the package iNEXT in R to standardize for varied sampling intensities (v3.0.2; Hsieh et al., 2025). We evaluated these variables as potential mechanisms for a “luxury effect” through which wealth index may influence arthropod community composition. For each relationship, we fit a Bayesian regression model using the R package brms (v2.23.0; Bürkner et al., 2025). Variables that were approximately normally distributed—garden floral richness and Shannon diversity, landscape tree canopy cover, and landscape floral richness and Shannon diversity— were modeled using Gaussian distributions. We used a lognormal model for garden floral species per m², which was right-skewed. For variables that were strictly positive, right-skewed, and with a high number of zero values, we used a hurdle-gamma distribution with a log link: local shrub and tree cover, bare ground, garden floral species/m², landscape water and wetland cover, and shrub and grassland cover. All models were fit using Hamiltonian Monte Carlo sampling with four chains and 6,000 total post-warmup draws. Convergence was assessed using R values and effective sample sizes, and model fit was evaluated via posterior predictive checks (Fig. S2.1 & S2.2). Selection of priors is discussed in Appendix S2: Supplementary methods and Table S2.1.

#### Community Dissimilarity

We tested whether local and landscape floral communities, arthropod functional group community composition, and bee communities differed across our sites, and whether these differences were associated with impervious surface and neighborhood wealth. We visualized each of our communities using Non-metric Multidimensional Scaling (NMDS) ordination with Bray-Curtis distances in the R package vegan (v2.7.3, Oksanen et al., 2026). We then analyzed the relationship between our continuous environmental covariates (impervious surface and wealth) and each community using a distance-based multivariate multiple regression permutation test with Bray-Curtis distances with the *adonis2* function in vegan. We also analyzed differences in our arthropod functional groups and bee community composition by month using a permutational multivariate analysis of variance with *adonis2*. For all analyses, significance was assessed using 999 permutations. To verify that significant PERMANOVA results reflected differences in community composition rather than differences in multivariate dispersion, we tested for homogeneity of group dispersions using the *betadisper* function followed by permutation tests (PCoA plot of group centroids with dispersion shown in Fig. S1.3). To visualize continuous environmental variables on ordination plots, we used the *ordisurf* function in vegan to fit generalized additive models (GAMs) onto the ordination space. This approach models smooth gradients of each variable across the two-dimensional ordination, which are displayed as contour lines. Finally, to see which species were driving differences in community composition, we ran multivariate generalized linear models (GLMs) in the R package mvabund, which is used for high-dimensional ecological abundance data and uses a negative binomial distribution by default (v4.2.8; Wang et al., 2025). The *mvabund* function fits generalized linear regressions to each species and utilizes resampling-based hypothesis testing to generate p-values.

#### Arthropod Abundance and Bee Diversity

We analyzed the relationship between wealth and impervious surface, and total arthropod abundance from the summer, using a Bayesian mixed-effects negative binomial regression in the brms package in R (v2.23.0; Bürkner et al., 2025). We included *Impervious Surface*, *Wealth Index,* and *Month* as fixed effects and *site* as a random effect. *Impervious Surface* and *Wealth Index* were standardized (mean = 0, SD = 1) to improve model performance and comparison across variables, such that coefficient estimates represent the effect of a one standard deviation change in each predictor. To select the spatial scale for impervious surface, we evaluated candidate models at buffer radii of 200 m and 500 m using leave-one-out cross-validation. Although both scales yielded comparable predictive accuracy, the 200 m model showed slightly superior out-of-sample performance (ΔELPD = 0.4). We retained the 200 m scale for final analyses, as smaller scales align with previously documented landscape effects on insect populations (Kaiser & Resasco, 2024). We also modeled pollinator, herbivore, and natural enemy abundance with the same model structure. To assess the impact of our environmental covariates on our bee biodiversity, we ran Bayesian mixed-effects regressions for three biodiversity indices — observed bee species richness, Shannon diversity, and Simpson diversity — which are sensitive to rare, intermediate, and dominant species, respectively. We did not rarefy our bee biodiversity estimates, since our sampling effort was standardized across sites. For each model, *Impervious Surface*, *Wealth Index,* and *Month* as fixed effects and *site* as a random effect. We added a spline-smoothing term to our Wealth Index predictor to address the hypothesized non-linearity in our response variable. Bee species richness was modeled with a Poisson distribution, and bee Shannon and Simpson diversity were modeled with a log-normal distribution. All models were fit using Hamiltonian Monte Carlo sampling with four chains and 6,000 total post-warmup draws. Convergence was assessed using R values and effective sample sizes, and model fit was evaluated via posterior predictive checks (Fig. S2.3). Selection of priors is discussed in Appendix 2: Supplementary methods and Table S2.2.

#### Structural Equation Path Analysis

We used a Bayesian structural equation modeling (SEM) framework to evaluate the direct and indirect pathways connecting landscape/socioeconomic drivers (impervious surface and neighborhood wealth) to crop output (Δ cucumber abundance, defined as the difference in fruit count between open-pollinated treatment and hand-pollinated control plants). We initially hypothesized pathways mediated by overall arthropod abundance and multiple diversity metrics (conceptual path diagram, Fig. S1.2a). However, to ensure model convergence and prevent overfitting, we streamlined the SEM framework based on preliminary exploratory regressions. Because preliminary univariate models showed no meaningful relationships between cucumber production and general pollinator, herbivore, or natural enemy abundances (Table S1.9), these variables were excluded from the final SEM structure.

Our finalized SEM comprised two chained submodels fit within the brms R package: 1) bee Shannon diversity as a function of impervious surface and wealth index, and 2) Δ cucumber abundance as a function of bee Shannon diversity, impervious surface, and wealth index (Fig. S1.2b). Bee Shannon diversity was scaled and modeled using a Gaussian distribution, while Δ cucumber abundance was modeled using a Student’s t distribution to accommodate potential outliers and non-normal error structures. All continuous predictors were standardized (mean = 0, SD = 1) prior to analysis to facilitate comparison of effect sizes and improve model performance. Models were fit using Hamiltonian Monte Carlo sampling with four chains and 6,000 total post-warmup draws. Convergence was assessed using R values and effective sample sizes, and model fit was evaluated via posterior predictive checks (Fig. S2.4 & S2.5). Selection of priors discussed in Appendix 2: Supplementary methods and Table S2.3.

## RESULTS

A total of 7000 arthropods were sampled across all community garden sites. Of this total, 1598 (22.8%) were categorized as pollinators, 2691 (38.4%) as herbivores, and 912 (13%) as natural enemies (all functional group classifications shown in Table S1.2). The most collected taxa were thrips (Thysanoptera, n = 2060), longhorn bees (*Melissodes*, n = 684; *Eucera*, n = 460), ants (Formicidae, n = 466), and minute pirate bugs (Anthocoridae; *Orius*, n = 402). A total of 191 cucumber fruits were harvested over the course of the summer across 20 sites. One site, Jefferson Green, did not produce cucumbers on any of the cucumber plants.

### Local and Landscape Variables and Wealth

We found little evidence that neighborhood wealth was associated with variation in either local garden characteristics or surrounding landscape composition (Table 1). None of the local variables showed a clear relationship with Wealth Index, as all 95% credible intervals overlapped zero. Effect sizes were generally small and uncertain: wood cover (β = 0.17, 95% CI: [−0.31, 0.69]), garden floral richness (β = 2.37, 95% CI: [−4.55, 9.17]), floral Shannon diversity (β = 2.73, 95% CI: [−2.63, 7.94]), bare ground (β = −0.55, 95% CI: [−1.12, 0.11]), and floral density (β = −0.07, 95% CI: [−0.28, 0.15]). Similarly, we observed no strong relationships between Wealth Index and landscape-scale variables within 500–1000 m of garden sites. Estimates for tree canopy cover (β = 1.69, 95% CI: [−0.21, 3.54]), landscape floral richness (β = −5.04, 95% CI: [− 20.10, 10.51]), floral Shannon diversity (β = −1.91, 95% CI: [−13.21, 8.97]), water and wetland cover (β = −0.03, 95% CI: [−0.52, 0.62]), and shrub and grassland cover (β = −0.14, 95% CI: [− 1.18, 0.84]) were all insignificant, with credible intervals overlapping zero.

**Table 1.** Bayesian posterior summaries for each local and landscape response variable as a function of wealth index across garden sites. Bolded values indicate significant effects where credible intervals do not overlap zero.

| Response Variable | Effect of 1 SD increase<br>in Wealth Index | SE | Lower 95% CI | Upper 95% CI |
| --- | --- | --- | --- | --- |
| **Local Variables** |  |  |  |  |
| Tree and shrub cover (m<sup>2</sup>) | 0.17 | 0.25 | -0.31 | 0.69 |
| Garden flora richness | 2.37 | 3.47 | -4.55 | 9.17 |
| Garden flora Shannon<br>diversity | 2.73 | 2.68 | -2.63 | 7.94 |
| Bare ground (m<sup>2</sup>) | -0.55 | 0.31 | -1.12 | 0.11 |
| Flora density (species/m<sup>2</sup>) | -0.07 | 0.11 | -0.28 | 0.15 |
| **Landscape Variables** |  |  |  |  |
| Tree canopy cover (500 m) | 1.69 | 0.93 | -0.21 | 3.54 |
| Landscape flora richness<br>(1000 m) | -5.04 | 7.72 | -20.10 | 10.51 |
| Landscape flora Shannon<br>diversity (1000 m) | -1.91 | 5.60 | -13.21 | 8.97 |
| Water & wetland (1000 m) | -0.03 | 0.29 | -0.52 | 0.62 |
| Shrub & grassland (1000 m) | -0.14 | 0.51 | -1.18 | 0.84 |

### Community Dissimilarity

Community composition responses to environmental and temporal variables varied across taxa (Table S1.4). Local floral community composition was not significantly associated with either impervious surface (R^2^ = 0.09, F = 1.89, p = 0.126, Fig. 3a) or wealth index (R^2^ = 0.05, F = 0.96, p = 0.39) whereas landscape floral community composition showed a modest but significant relationship with impervious surface (R^2^ = 0.10, F = 2.09, p = 0.030, Fig. 3b), but not wealth index (R^2^ = 0.05, F = 1.03, p = 0.31). In contrast, arthropod and bee communities exhibited strong temporal turnover, with month explaining substantial variation in both arthropod functional group composition (R^2^ = 0.26, F = 10.72, p = 0.001, Fig. 3e) and bee species composition (R^2^ = 0.29, F = 10.69, p = 0.001, Fig. 3f). Dispersion did not differ among months for arthropod functional groups (p = 0.84, Fig. S1.3a, Table S1.5) but differed significantly for bee species communities (p < 0.01, Fig 1.3b, Table S1.5). Across the full summer period, impervious surface was a significant predictor of both arthropod functional group composition (R^2^ = 0.26, F = 6.73, p = 0.001, Fig. 3c) and bee species composition (R^2^ = 0.14, F = 3.44, p = 0.026, Fig. 3d), whereas wealth index had no effect on arthropod functional groups but was significantly associated with bee species composition (R^2^ = 0.10, F = 2.36, p = 0.039).

**Figure 3.**
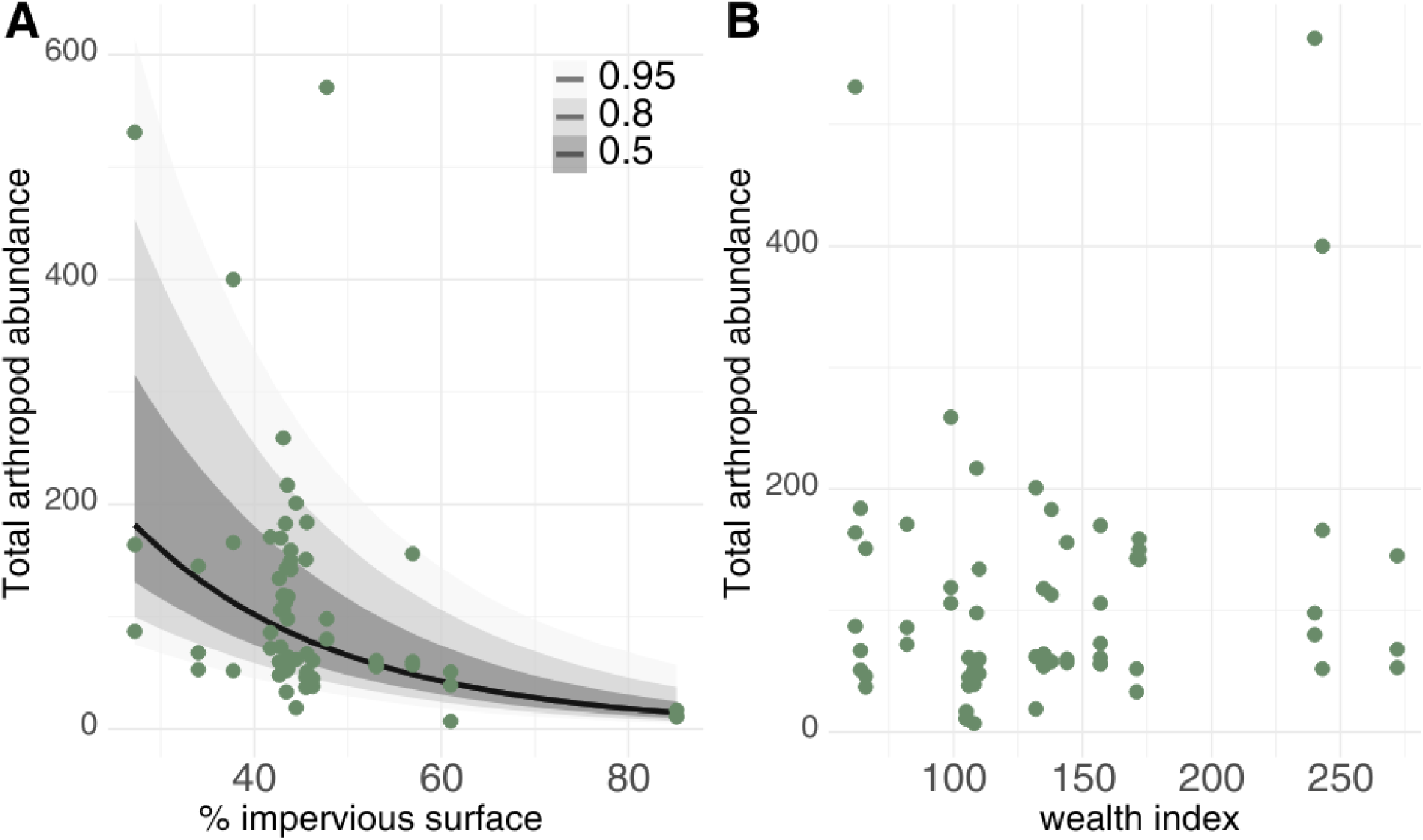
Scatterplots of total arthropod abundance and **a)** impervious surface at 200 meters (β = −0.50, 95% CI: [−0.68, −0.31]), with 50, 80, and 95% credible intervals and **b)** wealth index at 500 meters plotted for the full summer data.

The results of our multivariate GLMs showed that several arthropod functional groups responded significantly to impervious surface (Deviance = 54.26, p = 0.001), with pollinators (p = 0.010), herbivores (p = 0.035), and predators (p < 0.01) driving the trend. The multivariate GLM for bee community composition indicated significant effects of impervious surface (Deviance = 48.24, p = 0.012), while wealth index had no significant effect (p = 0.277). These results suggest that observed community-level differences may be driven by diffuse shifts across multiple species rather than strong responses of individual taxa. The only species related to wealth index was *Apis mellifera,* the western honeybee, whose abundance increased by 1.9% for every unit increase in wealth index (β = 0.019, p = 0.039).

### Arthropod Abundance

Total arthropod abundance strongly varied with impervious surface, with abundance decreasing by 39% for every standard deviation increase in impervious surface values (β = −0.50, 95% CI: [− 0.68, −0.31], Table S1.6). All three focal arthropod functional groups (pollinators, herbivores, and natural enemies) decreased significantly as impervious surface increased. Pollinators decreased by 39.9% (β = −0.51, 95% CI: [−0.78, −0.24]), herbivores by 36.4% (β = −0.45, 95% CI: [−0.75, −0.13]), and natural enemies by 33.9% (β = −0.41, 95% CI: [−0.60, −0.23]) for every standard deviation increase in impervious surface (Fig 4, Table S1.7). No group abundances varied significantly with wealth index (Table S1.7).

**Figure 4.**
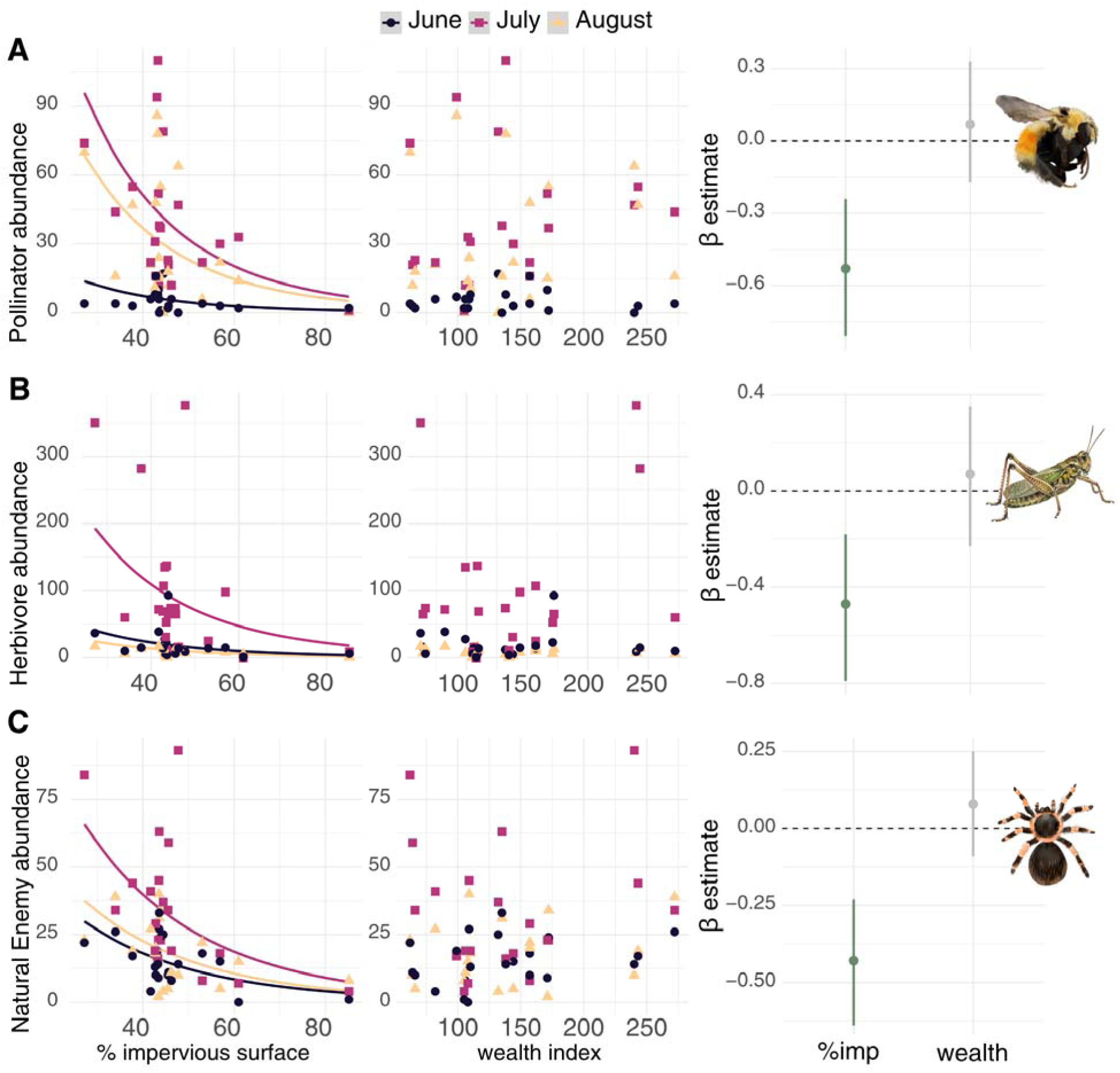
Relationships between arthropod functional group abundance and environmental gradients across sites. Panels show responses of **a)** pollinators, **b)** herbivores, and **c)** natural enemies to percent impervious surface and wealth index, with Bayesian GLM predictions overlaid by month. The right panels display model slope coefficients with 95 % credible intervals. All three groups declined significantly as impervious surface increased. No groups showed a significant association with wealth. *Image credits*: a) Asia Kaiser, b-c) Canva Pro License.

**Figure 5.**
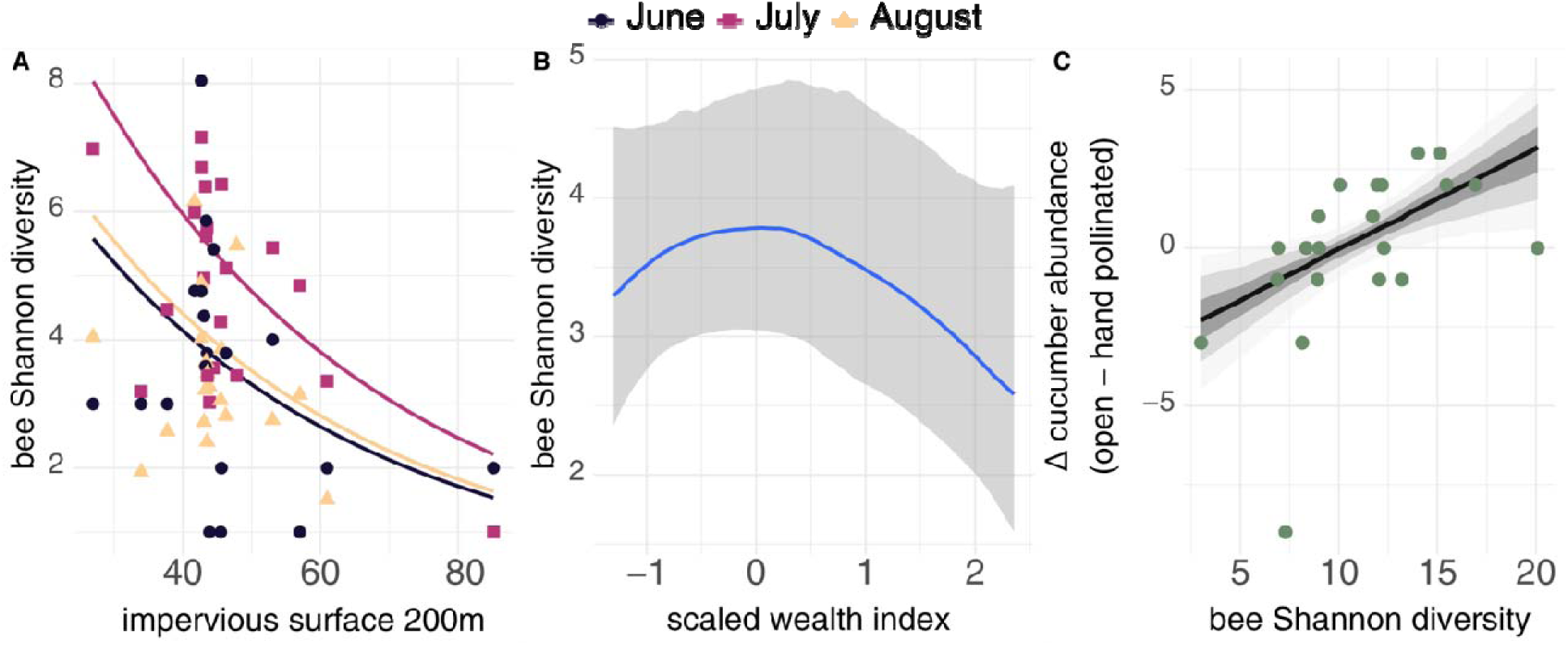
Relationships between bee Shannon diversity and **a)** percent impervious surface at 200 meters and **b)** conditional effects of neighborhood wealth index (scaled) modeled with a smoothing spline. Points represent site-level observations, with Bayesian GLM predictions shown by month. Bee Shannon diversity declined significantly with increasing impervious surface (β = −0.25, 95% CI: [−0.40, −0.11]) and showed a hump-shaped response to wealth (spline SD = 0.16, 95% CI: [0.01, 0.53]), increasing at low to moderate wealth levels and decreasing at higher wealth levels. Panel **c)** shows the relationship between bee Shannon diversity and the difference in cucumber production between open-pollinated (treatment) and hand-pollinated (control) plants. Higher Shannon diversity was associated with greater cucumber production under open pollination relative to hand pollination (β = 1.18, 95% CI: [0.27, 2.09]). Shaded bands represent the 50%, 80%, and 95% credible intervals.

### Bee Biodiversity

Bee species richness, Shannon diversity, and Simpson diversity were all strongly associated with surrounding impervious surface cover (Table S1.8). For every one standard deviation increase in impervious surface, there was an estimated decrease of approximately 0.71 bee species (β = − 0.34, 95% CI [−0.51, −0.17]), and Shannon diversity and Simpson diversity decreased by 0.25 (β = −0.25, 95% CI [−0.40, −0.11]) and 0.20 (β = −0.20, 95% CI [−0.34, −0.05]) units, respectively.

We observed a non-linear relationship between wealth and bee species richness (spline SD = 0.20, 95% CI [0.02, 0.60]) and Shannon diversity (spline SD = 0.16, 95% CI [0.01, 0.53]), with outcomes peaking at intermediate wealth values (Fig. S1.4, Table S1.8). Extracting peak diversity estimates from posterior draws of our models revealed that Shannon diversity peaked at a wealth index of 135 and richness at 146, representing wealth levels 35% and 46% higher than the US average of 100. While Simpson diversity showed a hump-shaped trend (Figure S1.4), the credible intervals for its smoothing parameter overlapped zero, suggesting an uncertain non-linear trend (spline SD = 0.13, 95% CI [0.00, 0.46]).

### Structural Equation Path Analysis

Δ cucumber abundance was positively associated with bee Shannon diversity (scaled bee Shannon diversity β = 1.18, 95% CI: [0.27, 2.09], Fig. 6). Cucumber abundance was the only cucumber outcome variable significantly related to Bee Shannon diversity (seed set, fruit mass, fruit abundance, and cylindricity outcomes shown in Fig. S1.5). In separate regressions examining individual arthropod groups, none of the other arthropod predictors—pollinator abundance, herbivore abundance, natural enemy abundance, or bee species richness—showed a clear relationship with cucumber production (Table S1.9). Based on these results, these paths were excluded from the structural equation model to reduce complexity, avoid overfitting, and improve convergence (Fig. S2.2b).

There was no clear relationship between impervious surface, wealth index, and cucumber production, with credible intervals overlapping zero. There was a statistically significant negative indirect effect of impervious surface on cucumber outcomes mediated through bee Shannon diversity (β = −0.59, 95% CI [−1.42, −0.07]). This indicates that higher impervious cover reduces bee diversity, which in turn leads to a measurable decrease in cucumber yields/set.

## DISCUSSION

The aim of this study was to investigate how landscape-level biophysical and socioeconomic features shape arthropod community composition at community garden sites and, subsequently, the ecosystem services provided, as measured by crop yield. We found that total arthropod abundance is strongly and negatively affected by surrounding impervious surfaces at garden sites, and that changes in pollinator, herbivore, and predator abundances are driving associated changes in community composition. Pollinators had the steepest decrease, followed by herbivores and natural enemies (predators, parasitoids, predator-omnivores). This suggests that pollinators may be especially sensitive to urbanization.

In addition, we found that bee species richness and Shannon diversity show a significant relationship with neighborhood wealth levels. We observed a nonlinear relationship between neighborhood wealth and bee richness and Shannon diversity, with the highest levels at wealth values between 35-46% above the national average (Fig. S1.4). One potential mechanism could be competition between native bees and managed honeybees, as our multivariate GLM of species responses to wealth showed *Apis mellifera,* the western honeybee, as the only species related to community dissimilarity by wealth. We observed that cucumber production was related to bee Shannon diversity, but not to species richness. This indicates that fruit production in this crop may be more sensitive to the evenness of bee communities than to the total number of species, as there needs to be enough effective pollinators. Increases in species richness, but not evenness, may be driven by rare taxa that do not substantially contribute to overall diversity or pollination function. We also found no independent effect of impervious surface or wealth on our cucumber abundance outcomes and demonstrated that bee diversity is the primary mediator of differences in fruit set in this system. This emphasizes that maintaining diverse wild bee assemblages in urban agroecosystems is one of the most important factors to ensure optimal crop yields.

The negative effects of impervious surface on total arthropod abundance and on our three focal groups (pollinators, herbivores, and natural enemies) support broader evidence that urbanization and habitat loss is a key driver of global insect declines (Piano et al., 2020; Wagner, 2020) and were consistent with our hypotheses and previous studies (Geslin et al., 2016; Kaiser & Resasco, 2024; Maher et al., 2022) (Fig. 4). In our system, impervious surface influenced not only overall abundance but also the functional composition of arthropod communities at garden sites (Fig. 3c). Maher et al. (2022) found similar uneven declines in arthropod biodiversity across taxa with increasing impervious surface, and these effects were mediated by local abiotic conditions such as humidity and temperature, which were not measured in our study. If arthropod groups differ in their sensitivity to these abiotic factors, such variation could drive shifts in community composition along impervious surface gradients, rather than produce uniform declines across all groups.

Unexpectedly, wealth index was not associated with local or landscape biotic variables, nor with most arthropod responses. This contrasts with studies documenting stronger luxury effects in arid urban systems, typically expressed as increased floral richness and tree cover with wealth (Chamberlain et al., 2020; Hope et al., 2003). However, evidence for luxury effects in arthropods remains understudied (Leong et al., 2018), and the high availability of habitat and resources in urban community gardens may buffer arthropod communities from socioeconomic-driven differences in matrix quality surrounding gardens. A meta-analysis of diversification practices in agroecosystems found that intercropping was most effective at reducing herbivore abundance while increasing natural enemies (Seimandi-Corda et al., 2026). Because all community gardens in our system already maintain diverse plantings with a variety of crops in close proximity and additional floral resources, the abundance of less-mobile beneficial arthropods may already be optimized in these systems. Thus, community gardens may help buffer arthropod communities against luxury effects across neighborhoods with varying socioeconomic legacies and wealth levels. Increasing ground-level habitat complexity by adding straw mulch or leaf litter has been shown to improve pest control and increase the abundance of natural enemies in these systems (Ploessl et al., 2024). This local management approach may be more effective at enhancing these ecosystem services than changing broader landscape features to benefit these arthropod groups.

Bee species richness was the only arthropod response associated with wealth, peaking at intermediate levels and declining in both low- and high-income neighborhoods, consistent with previous work showing bee biodiversity is lower in very high-income neighborhoods (Kaiser & Resasco, 2024). The mechanism remains unclear, as wealth was unrelated to local habitat (floral diversity, vegetation cover, bare ground) or landscape context (tree canopy, floral diversity, wetland, shrub, and grassland cover) that would indicate differences in habitat quality. This pattern may instead reflect unobserved landscape-level processes. For example, impervious surface has been shown to structure bee assemblages independently of local floral composition (Geslin et al., 2016) and high-income neighborhoods often feature distinct management regimes. Specifically, affluent areas may harbor more managed honeybee colonies, which have been shown to outcompete wild bees in urban areas (MacInnis et al., 2023). Wealthy neighborhoods may also have higher residential pesticide use, which has been shown to have sublethal effects on bees in urban environments, reducing their fitness (Siviter et al., 2023). Given their larger foraging ranges than other arthropod groups, bees may be more responsive to these broader landscape effects (Kremen et al., 2007).

To assess arthropod-mediated ecosystem services, we selected cucumber (*Cucumis sativus*) as a single focal sentinel crop due to its monoecious flowering strategy and strong reliance on insect pollination. However, because herbivore control (Motzke et al., 2015) and leaf damage (Barber et al., 2011) exert minimal influence on cucumber yield compared to pollination (Motzke et al., 2015), future research should incorporate multiple crop species to evaluate how simultaneous arthropod drivers interact across diverse crop traits. Additionally, while using potted plants standardized water and soil conditions across garden sites, this approach isolated crops from belowground interactions. Given that root herbivory can substantially suppress cucumber yield (Barber et al., 2011), in-ground sentinel plantings will be essential for future studies to capture the full impact of soil-dwelling arthropods.

We utilized passive arthropod sampling methods to allow simultaneous sampling across 21 study sites. However, passive methods may underrepresent key taxa that disproportionately affect cucumber plants. Combining passive traps with active sampling (e.g., hand-netting) in future work will better resolve specific plant–arthropod interactions and capture taxa missed by passive traps (Gibbs et al., 2017; Prendergast et al., 2020). In addition, we categorized arthropod functional groups by the life stage they were at when collected. Most of our arthropods, however, were in the adult life stage, and thus we may be under-sampling juveniles with different functional feeding strategies, biasing our estimates of the overall functional-group community composition in our gardens. Still, we believe our results remain informative, as the objective of our study was to compare sites across an urbanization and wealth gradient to identify the landscape drivers of arthropod-mediated ecosystem services, rather than to fully characterize arthropod community composition at each site.

## CONCLUSION

As urban land use and urban populations grow globally, understanding the factors shaping arthropod biodiversity in these landscapes is increasingly important for conserving biodiversity and the ecosystem services these arthropods provide. Our results show that impervious surfaces in cities strongly affect not only arthropod abundance but also the relative abundance of different functional groups and community composition. In addition, we observe a luxury effect: socioeconomic factors correlate with bee richness and diversity in cities, but the relationship is non-linear, peaking at moderate wealth levels. Unexpectedly, wealth did not correlate with any variables related to habitat quality at sites or in the surrounding matrix, indicating the mechanisms driving this pattern warrant further investigation. The influence of non-native western honeybees, pesticide use, and lawn water use could be driving this decline in bee biodiversity in high-wealth areas and should be examined in future studies.

## ACKNOWLEDGMENTS

We thank the members of the Resasco Lab for feedback on the manuscript and Rhiannon Danborn for her assistance with fieldwork and data collection. We also thank Denver Urban Gardens for their support and partnership in this research project. This research was supported by a USDA NIFA predoctoral fellowship (grant COLW-2023-11576).

## AUTHOR CONTRIBUTIONS

A.K. and J.R. conceived the study; N.M. and R.A provided further input on the study design. All authors assisted with data collection. A. K. performed the data analyses and wrote the first draft of the manuscript. J.R., N.M, and R.A. reviewed and provided feedback on subsequent drafts. All authors have read and approved the final manuscript.

## COMPETING INTERESTS

The authors declare no competing interests.

## Notes

https://github.com/asiakaiser/dug-research

### Competing Interest Statement

The authors have declared no competing interest.

