## Supplemental Materials for "Urbanization and wealth alter arthropod biodiversity and pollination services in community gardens"

**Open Research Statement:** The data and statistical code that support the findings of this study are openly available at the GitHub repository: <https://github.com/asiakaiser/dug-research>.

**Appendix S1: Extended Results & Summary Outputs**

**Table S1.1 Summary of community garden site characteristics.** Site name, Garden size (m²), number of plots, 2025 ESRI Wealth Index (within a 500 m radius), and percent impervious surface and tree canopy cover (within a 200 m radius) for community gardens in the Denver metropolitan area, Colorado, USA (<https://dug.org/>).

|  | Site | Garden Size (m^2^) | No. of Garden Plots | Wealth Index | % Impervious surface | % Tree Canopy Cover |
| --- | --- | --- | --- | --- | --- | --- |
| 1) | Applewood | 627 | 28 | 240 | 47.74 | 10.23 |
| 2) | Charles Hay | 459 | 25 | 99 | 43.09 | 15.27 |
| 3) | Eiber School | 579 | 24 | 109 | 43.53 | 14.19 |
| 4) | Gables School | 734 | 32 | 171 | 43.41 | 14.89 |
| 5) | Samuels School | 1642 | 55 | 157 | 42.82 | 16.24 |
| 6) | Slater Elementary School | 635 | 26 | 110 | 42.67 | 13.62 |
| 7) | West Colfax | 1108 | 38 | 82 | 41.72 | 12.36 |
| 8) | Greenway | 776 | 35 | 144 | 56.92 | 7.97 |
| 9) | Cook Park | 1175 | 48 | 172 | 43.89 | 13.33 |
| 10) | Gabriel Cam Memorial | 738 | 30 | 64 | 45.61 | 9.75 |
| 11) | KCAA | 332 | 18 | 106 | 46.25 | 14.04 |
| 12) | Park Hill School | 738 | 16 | 243 | 37.74 | 17.72 |
| 13) | Ruby Hill Park | 858 | 56 | 62 | 27.20 | 10.23 |
| 14) | Sabin School | 887 | 27 | 135 | 43.59 | 14.73 |
| 15) | Shoshone | 415 | 12 | 108 | 60.99 | 9.58 |
| 16) | Palmer School | 649 | 20 | 138 | 43.31 | 14.22 |
| 17) | Edison School | 523 | 21 | 132 | 44.45 | 16.23 |
| 18) | Market Station | 489 | 33 | 105 | 85.16 | 0.58 |
| 19) | Steck School | 465 | 17 | 272 | 34.00 | 20.82 |
| 20) | Jefferson Green | 1104 | 28 | 66 | 45.51 | 12.30 |
| 21) | GW School | 474 | 13 | 157 | 53.04 | 7.10 |

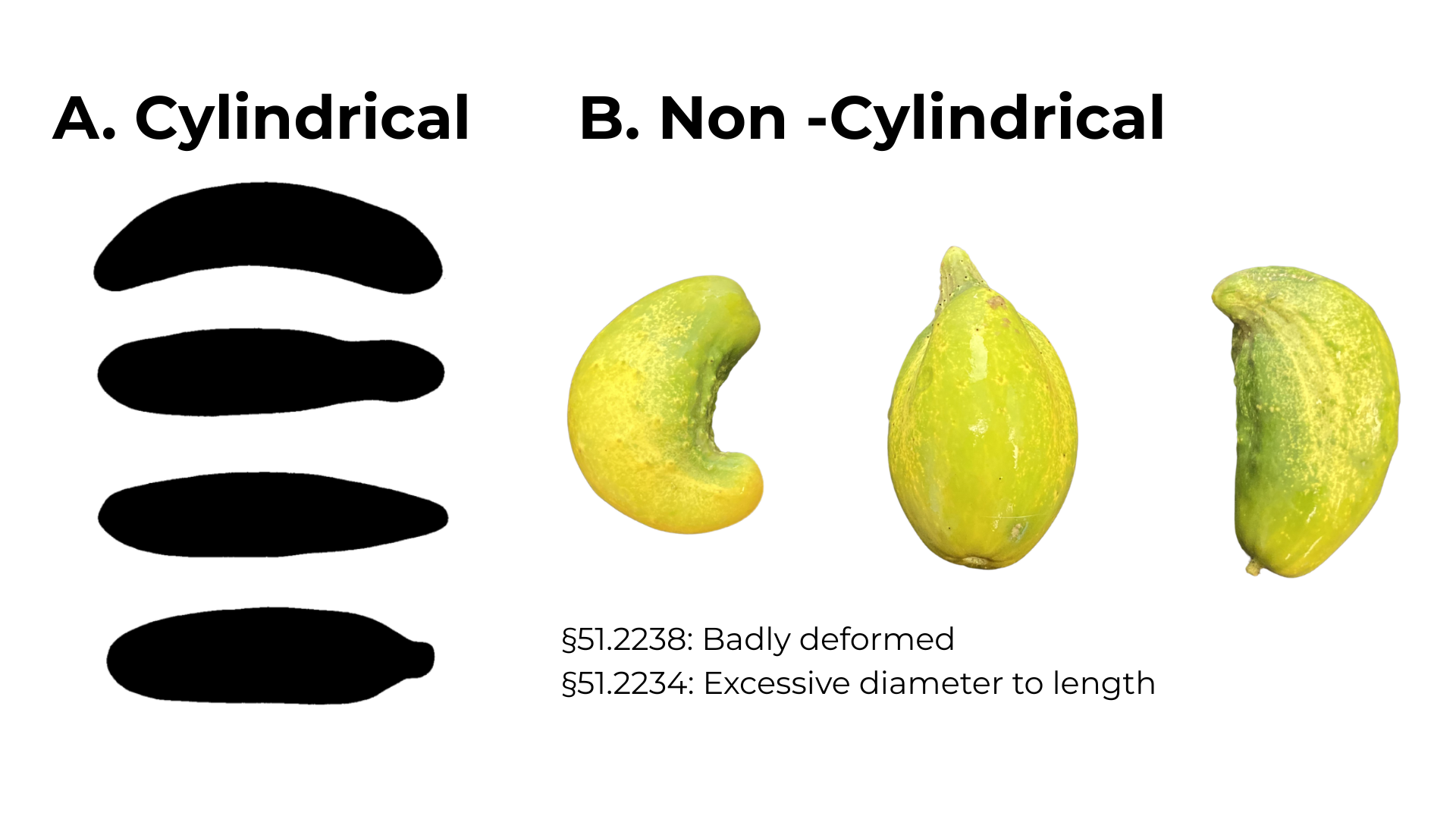

**Figure S1.1 Cucumber cylindricity.** Examples of **a)** cylindrical and **b)** non-cylindrical cucumber classifications with section statute numbers from the United States Standards for Grades of Cucumbers (USDA, 2016). Cylindrical examples are the “minimum shapes permissible in U.S. No. 1 Grade”. Photo credits: **a)** United States Department of Agriculture (USDA), **b)** Asia Kaiser.

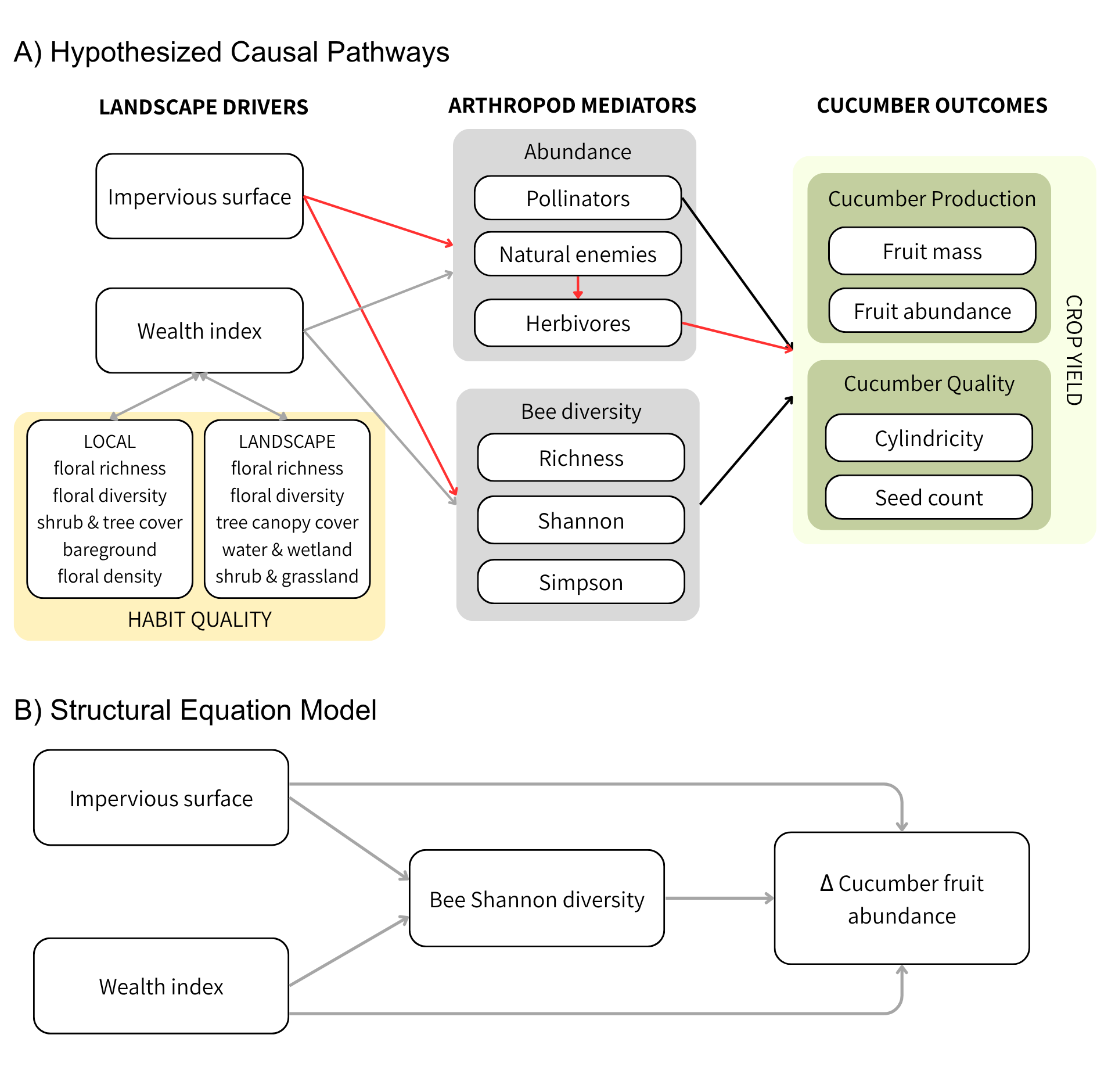
**Figure S1.2** Hypothesized causal framework and final structural model structure. **A)** Directed acyclic graph representing hypothesized relationships among landscape environmental factors (including local and landscape habit quality), socioeconomic drivers, arthropod communities, and cucumber outcomes. **B)** Diagram of the final Structural Equation Model (SEM) structure, testing the direct and indirect effects of impervious surface cover and wealth index on Δ cucumber fruit abundance as mediated by bee Shannon diversity. Nonsignificant mediating variables are omitted. Line colors indicate relationship types: gray lines represent non-linear relationships or unknown directions; black lines indicate positive relationships; red lines indicate negative relationships.

**Table S1.2 Denver urban gardens arthropod community.** The taxonomic identity, abundance, and functional group designation of all arthropods collected during this study (June - August 2025). All specimens were identified to the lowest taxonomic level needed to determine feeding behavior.

| **Class** | **Order** | **Family** | **Genus** | **Functional Group** | **Abundance** |
| --- | --- | --- | --- | --- | --- |
| Arachnida | Acariformes | Unknown | Unknown | Parasite | 2 |
|  |  |  | Unknown | Unknown | 147 |
|  | Araneae | Dysderidae | Dysdera | Predator | 9 |
|  |  | Linyphiidae | Unknown | Predator | 1 |
|  |  | Lycosidae | Schizocosa | Predator | 2 |
|  |  | Salticidae | Unknown | Predator | 10 |
|  |  | Thomisidae | Unknown | Predator | 1 |
|  |  | Unknown | Unknown | Predator | 152 |
|  | Ixodida | Ixodidae | Unknown | Sanguivore | 1 |
|  |  | Unknown | Unknown | Sanguivore | 1 |
|  | Opiliones | Unknown | Unknown | Predator | 4 |
|  |  |  |  | Predator/Scavenger | 50 |
|  | Pseudoscorpiones | Unknown | Unknown | Predator | 3 |
|  | Trombidiformes | Anystidae | Unknown | Predator | 1 |
|  |  | Tetranychidae | Tetranychus | Herbivore | 1 |
|  |  | Unknown | Unknown | Herbivore | 12 |
|  |  |  | Unknown | Unknown | 12 |
|  | Unknown | Unknown | Unknown | Predator | 10 |
|  |  |  |  | Unknown | 26 |
| Chilopoda | Unknown | Unknown | Unknown | Predator | 24 |
| Clitellata | Unknown | Unknown | Unknown | Detritivore | 1 |
| Collembola | Unknown | Unknown | Unknown | Detritivore | 90 |
| Diplopoda | Polyxenida | Unknown | Unknown | Detritivore | 1 |
|  | Unknown | Unknown | Unknown | Detritivore | 55 |
| Entognatha | Diplura | Unknown | Unknown | Predator | 2 |
|  | Protura | Unknown | Unknown | Detritivore | 2 |
| Gastropoda | Unknown | Unknown | Unknown | Herbivore | 1 |
| Insecta | Araneae | Unknown | Unknown | Predator | 1 |
|  | Coleoptera | Anthicidae | Anthicus | Scavenger | 1 |
|  |  | Anthocoridae | Orius | Predator | 1 |
|  |  | Carabidae | Bradycellus | Predator | 1 |
|  |  | Carabidae | Unknown | Predator | 5 |
|  |  | Cerambycidae | Unknown | Herbivore | 3 |
|  |  | Chrysomelidae | Unknown | Herbivore | 7 |
|  |  | Cicadellidae | Unknown | Herbivore | 2 |
|  |  | Cleridae | Chariessa | Predator | 1 |
|  |  | Coccinellidae | Coccinella | Predator | 1 |
|  |  |  | Unknown | Predator | 18 |
|  |  | Curculionidae | Scolytus | Herbivore | 13 |
|  |  |  | Sphenophorus | Herbivore | 2 |
|  |  |  | Unknown | Herbivore | 22 |
|  |  |  | Xyleborinus | Herbivore | 2 |
|  |  |  | Xylosandrus | Herbivore | 1 |
|  |  | Dermestidae | Unknown | Detritivore | 2 |
|  |  | Elateridae | Unknown | Herbivore | 5 |
|  |  | Erotylidae | Cryptophilus | Fungivore | 1 |
|  |  |  | Unknown | Fungivore | 1 |
|  |  | Lampyridae | Unknown | Predator | 4 |
|  |  | Latridiidae | Unknown | Fungivore | 7 |
|  |  | Meloidae | Epicauta | Herbivore | 7 |
|  |  |  | Unknown | Herbivore | 16 |
|  |  | Mordellidae | Unknown | Pollinator | 1 |
|  |  | Mycetophagidae | Unknown | Fungivore | 1 |
|  |  | Nitidulidae | Glischrochilus | Herbivore | 2 |
|  |  |  | Nitops | Detritivore | 25 |
|  |  |  |  | Herbivore | 15 |
|  |  | Scarabaeidae | Ataenius | Detritivore | 1 |
|  |  |  | Euphoria | Detritivore | 133 |
|  |  |  | Pleurophorus | Detritivore | 1 |
|  |  |  | Popillia | Herbivore | 81 |
|  |  |  | Unknown | Detritivore | 1 |
|  |  |  |  | Herbivore | 11 |
|  |  | Staphylinidae | Heterosilpha | Omnivore | 1 |
|  |  |  | Unknown | Omnivore | 1 |
|  |  |  |  | Predator | 29 |
|  |  | Tenebrionidae | Bothrotes | Detritivore | 1 |
|  |  |  | Unknown | Detritivore | 1 |
|  |  | Unknown | Unknown | Herbivore | 5 |
|  |  |  |  | Predator | 2 |
|  |  |  |  | Unknown | 168 |
|  | Dermaptera | Forficulidae | Forficula | Omnivore | 3 |
|  |  | Unknown | Unknown | Omnivore | 18 |
|  |  |  |  | Predator | 2 |
|  | Diptera | Anisopodidae | Sylvicola | Detritivore | 1 |
|  |  | Anthomyiidae | Delia | Herbivore | 2 |
|  |  |  | Unknown | Unknown | 1 |
|  |  | Calliphoridae | Protophormia | Scavenger | 1 |
|  |  |  | Unknown | Scavenger | 2 |
|  |  | Ceratopogonidae | Culicoides | Sanguivore | 1 |
|  |  |  | Unknown | Sanguivore | 1 |
|  |  | Chironomidae | Unknown | Detritivore | 5 |
|  |  | Chloropidae | Meromyza | Herbivore | 1 |
|  |  | Conopidae | Zodion | Parasitoid | 1 |
|  |  | Culicidae | Unknown | Sanguivore | 26 |
|  |  | Drosophilidae | Drosophila | Detritivore | 1 |
|  |  |  | Unknown | Detritivore | 1 |
|  |  | Muscidae | Musca | Detritivore | 4 |
|  |  |  | Unknown | Detritivore | 19 |
|  |  |  |  | Unknown | 18 |
|  |  | Mycetophilidae | Unknown | Fungivore | 1 |
|  |  | Polleniidae | Pollenia | Herbivore | 2 |
|  |  | Syrphidae | Toxomerus | Pollinator | 2 |
|  |  |  | Unknown | Pollinator | 3 |
|  |  | Tachinidae | Unknown | Parasitoid | 6 |
|  |  | Therevidae | Unknown | Predator | 1 |
|  |  | Unknown | Unknown | Detritivore | 4 |
|  |  |  |  | Herbivore | 1 |
|  |  |  |  | Pollinator | 1 |
|  |  |  |  | Sanguivore | 4 |
|  |  |  |  | Unknown | 188 |
|  | Hemiptera | Anthocoridae | Orius | Omnivore | 1 |
|  |  |  |  | Predator | 401 |
|  |  | Aphididae | Unknown | Herbivore | 133 |
|  |  |  |  | Herbivore | 1 |
|  |  | Cicadellidae | Agallia | Herbivore | 2 |
|  |  |  | Anascopus | Herbivore | 1 |
|  |  |  | Ceratagallia | Herbivore | 5 |
|  |  |  | Unknown | Herbivore | 200 |
|  |  | Cydnidae | Amnestus | Herbivore | 1 |
|  |  | Geocoridae | Geocoris | Predator | 3 |
|  |  | Miridae | Ceratocapsus | Herbivore | 2 |
|  |  |  | Chlamydatus | Herbivore | 3 |
|  |  |  |  | Predator | 1 |
|  |  |  | Lygus | Herbivore | 5 |
|  |  |  | Orthops | Herbivore | 3 |
|  |  |  | Pseudatomoscelis | Herbivore | 37 |
|  |  |  | Unknown | Herbivore | 2 |
|  |  |  |  | Predator | 1 |
|  |  | Nabidae | Nabis | Predator | 2 |
|  |  | Notonectidae | Buenoa | Predator | 1 |
|  |  | Pentatomidae | Thyanta | Herbivore | 1 |
|  |  | Reduviidae | Unknown | Predator | 4 |
|  |  | Rhopalidae | Brachycarenus | Herbivore | 1 |
|  |  | Rhyparochromidae | Emblethis | Detritivore | 1 |
|  |  |  | Megalonotus | Granivore | 1 |
|  |  |  |  | Herbivore | 1 |
|  |  | Tingidae | Unknown | Herbivore | 9 |
|  |  | Unknown | Unknown | Detritivore | 1 |
|  |  |  |  | Herbivore | 10 |
|  |  |  |  | Unknown | 100 |
|  | Hymenoptera | Andrenidae | Perdita | Pollinator | 4 |
|  |  |  | Protandrena | Pollinator | 1 |
|  |  | Apidae | Anthophora | Pollinator | 1 |
|  |  |  | Apis | Pollinator | 33 |
|  |  |  | Bombus | Pollinator | 43 |
|  |  |  | Brachymelecta | Kleptoparasite | 1 |
|  |  |  | Ceratina | Pollinator | 1 |
|  |  |  | Diadasia | Pollinator | 9 |
|  |  |  | Eucera | Pollinator | 460 |
|  |  |  | Melissodes | Pollinator | 688 |
|  |  |  | Melitoma | Pollinator | 4 |
|  |  |  | Svastra | Pollinator | 1 |
|  |  |  | Unknown | Pollinator | 11 |
|  |  | Bethylidae | Unknown | Parasitoid | 1 |
|  |  | Braconidae | Unknown | Parasitoid | 4 |
|  |  | Chalcididae | Unknown | Parasitoid | 6 |
|  |  | Eulophidae | Unknown | Parasitoid | 1 |
|  |  | Formicidae | Lasius | Omnivore | 1 |
|  |  |  | Tetramorium | Omnivore | 28 |
|  |  |  | Unknown | Omnivore | 446 |
|  |  | Halictidae | Agapostemon | Pollinator | 41 |
|  |  |  | Augochlorella | Pollinator | 8 |
|  |  |  | Halictus | Pollinator | 171 |
|  |  |  | Lasioglossum Dialictus | Pollinator | 64 |
|  |  |  | Lasioglossum Evylaeus | Pollinator | 14 |
|  |  |  | Lasioglossum Evylaeus | Unknown | 1 |
|  |  |  | Lasioglossum Sensu Stricto | Pollinator | 6 |
|  |  |  | Sphecodes | Kleptoparasite | 4 |
|  |  |  | Unknown | Pollinator | 2 |
|  |  | Megachilidae | Hoplitis | Pollinator | 1 |
|  |  |  | Lithurgopsis | Pollinator | 2 |
|  |  |  | Megachile | Pollinator | 5 |
|  |  |  | Osmia | Pollinator | 2 |
|  |  |  | Unknown | Pollinator | 1 |
|  |  | Mymaridae | Unknown | Parasitoid | 3 |
|  |  | Pompilidae | Unknown | Predator | 1 |
|  |  | Scelionidae | Unknown | Parasitoid | 2 |
|  |  | Sphecidae | Unknown | Parasitoid | 1 |
|  |  | Unknown | Unknown | Parasitoid | 17 |
|  |  |  |  | Pollinator | 4 |
|  |  |  |  | Predator | 8 |
|  |  | Vespidae | Polistes | Predator | 1 |
|  |  |  | Unknown | Predator | 26 |
|  |  |  | Vespula | Predator | 1 |
|  | Lepidoptera | Hesperiidae | Unknown | Pollinator | 3 |
|  |  | Papilionidae | Papilio | Pollinator | 1 |
|  |  | Unknown | Unknown | Pollinator | 8 |
|  |  |  |  | Unknown | 1 |
|  | Neuroptera | Chrysopidae | Unknown | Predator | 80 |
|  |  | Hemerobiidae | Unknown | Predator | 2 |
|  | Orthoptera | Acrididae | Unknown | Herbivore | 5 |
|  |  | Unknown | Unknown | Herbivore | 1 |
|  | Psocodea | Lachesillidae | Lachesilla | Detritivore | 3 |
|  | Thysanoptera | Aeolothripidae | Aeolothrips | Predator | 3 |
|  |  |  | Unknown | Predator | 1 |
|  |  | Thripidae | Unknown | Herbivore | 588 |
|  |  | Unknown | Unknown | Herbivore | 1467 |
|  |  |  |  | Predator | 1 |
|  | Trichoptera | Unknown | Unknown | Aquatic Scavenger | 2 |
|  | Unknown | Unknown | Unknown | Unknown | 22 |
| Malacostraca | Isopoda | Armadillidiidae | Armadillidium | Detritivore | 50 |
|  |  |  | Unknown | Detritivore | 104 |
|  |  | Unknown | Unknown | Detritivore | 43 |
| Unknown | Unknown | Unknown | Unknown | Unknown | 10 |

**Table S1.3. Denver urban gardens** **bee community.** The taxonomic identity and abundance of all bee species collected during this study (June - August 2025). Each individual was identified to the lowest possible taxonomic level.

| **Family** | **Genus** | **Specie** | **Abundance** |
| --- | --- | --- | --- |
| Andrenidae | Perdita | sp1 | 1 |
|  |  | sp2 | 1 |
|  |  | sp3 | 1 |
|  |  | spp. | 1 |
|  | Protandrena | spp. | 1 |
| Apidae | Anthophora | montana | 1 |
|  | Apis | mellifera | 33 |
|  | Bombus | bifarius | 3 |
|  |  | fervidus | 26 |
|  |  | huntii | 12 |
|  |  | nevadensis | 1 |
|  |  | pensylvanicus | 1 |
|  | Brachymelecta | interrupta | 1 |
|  | Ceratina | spp. | 1 |
|  | Diadasia | australis | 4 |
|  |  | enavata | 3 |
|  |  | rinconis | 2 |
|  | Eucera | pruinosa | 460 |
|  | Melissodes | agilis | 604 |
|  |  | bimaculatus | 66 |
|  |  | communis | 1 |
|  |  | coreopsis | 1 |
|  |  | niveus | 7 |
|  |  | sp1 | 1 |
|  |  | sp2 | 2 |
|  |  | spp. | 4 |
|  |  | tristis | 2 |
|  | Melitoma | grisella | 4 |
|  | Svastra | obliqua | 1 |
|  | Unknown | spp. | 11 |
| Halictidae | Agapostemon | angelicus/texanus | 11 |
|  |  | femoratus | 8 |
|  |  | virescens | 22 |
|  |  | aurata | 8 |
|  | Halictus | confusus | 1 |
|  |  | farinosus | 1 |
|  |  | ligatus | 10 |
|  |  | rubicundus | 105 |
|  |  | spp. | 1 |
|  |  | tripartitus | 53 |
|  | Lasioglossum Dialictus | semicaeruleum | 29 |
|  |  | spp. | 5 |
|  |  | versans | 1 |
|  |  | zephyrus | 29 |
|  | Lasioglossum Evylaeus | spp. | 15 |
|  | Lasioglossum Sensu Stricto | sisymbri | 1 |
|  |  | spp. | 5 |
|  | Sphecodes | davisii | 1 |
|  |  | spp. | 3 |
|  | Unknown | spp. | 2 |
| Megachilidae | Hoplitis | producta | 1 |
|  | Lithurgopsis | apicalis | 2 |
|  | Megachile | brevis | 1 |
|  |  | centuncularis | 1 |
|  |  | nivalis | 1 |
|  |  | parallela | 1 |
|  |  | rotundata | 1 |
|  | Osmia | inermis | 2 |
|  | Unknown | spp. | 1 |

**Table S1.4** **Community PERMANOVA model outputs.** Permutational Analysis of Variance (PERMANOVA) outputs for local and landscape floral families, arthropod functional groups, and bee species at each community garden site.

| **Community** | **Predictor** | **Df** | **R^2^** | **F value** | **P value** |
| --- | --- | --- | --- | --- | --- |
| Local Flora | Impervious surface | 1 | 0.091 | 1.894 | 0.126 |
|  | Wealth Index | 1 | 0.046 | 0.958 | 0.394 |
| Landscape Flora | Impervious surface | 1 | 0.100 | 2.094 | **0.030*** |
|  | Wealth Index | 1 | 0.048 | 1.034 | 0.314 |
| Monthly samples  Arthropod Functional Groups | Month | 2 | 0.263 | 10.7 | **0.001***** |
| Full summer  Arthropod Functional Groups | Impervious surface | 1 | 0.257 | 6.731 | **0.001***** |
|  | Wealth Index | 1 | 0.036 | 0.933 | 0.441 |
| Monthly samples  Bee species | Month | 2 | 0.287 | 10.69 | **0.001***** |
| Full summer  Bee species | Impervious surface | 1 | 0.138 | 3.435 | **0.026*** |
|  | Wealth Index | 1 | 0.095 | 2.361 | **0.039*** |

*P-values denoted by < .05*, < .01**, < .001****

**Table S1.5 Community dispersion test outputs.** Results of tests for homogeneity of multivariate dispersion used to assess assumptions of PERMANOVA models examining arthropod functional group and bee species community composition.

| **Community** | **Predictor** | **F value** | **P value** |
| --- | --- | --- | --- |
| Monthly samples - Arthropod Functional Groups | month | 0.174 | 0.845 |
| Monthly samples - Bee species | month | 5.731 | **< 0.01 **** |

*P-values denoted by < .05*, < .01**, < .001****

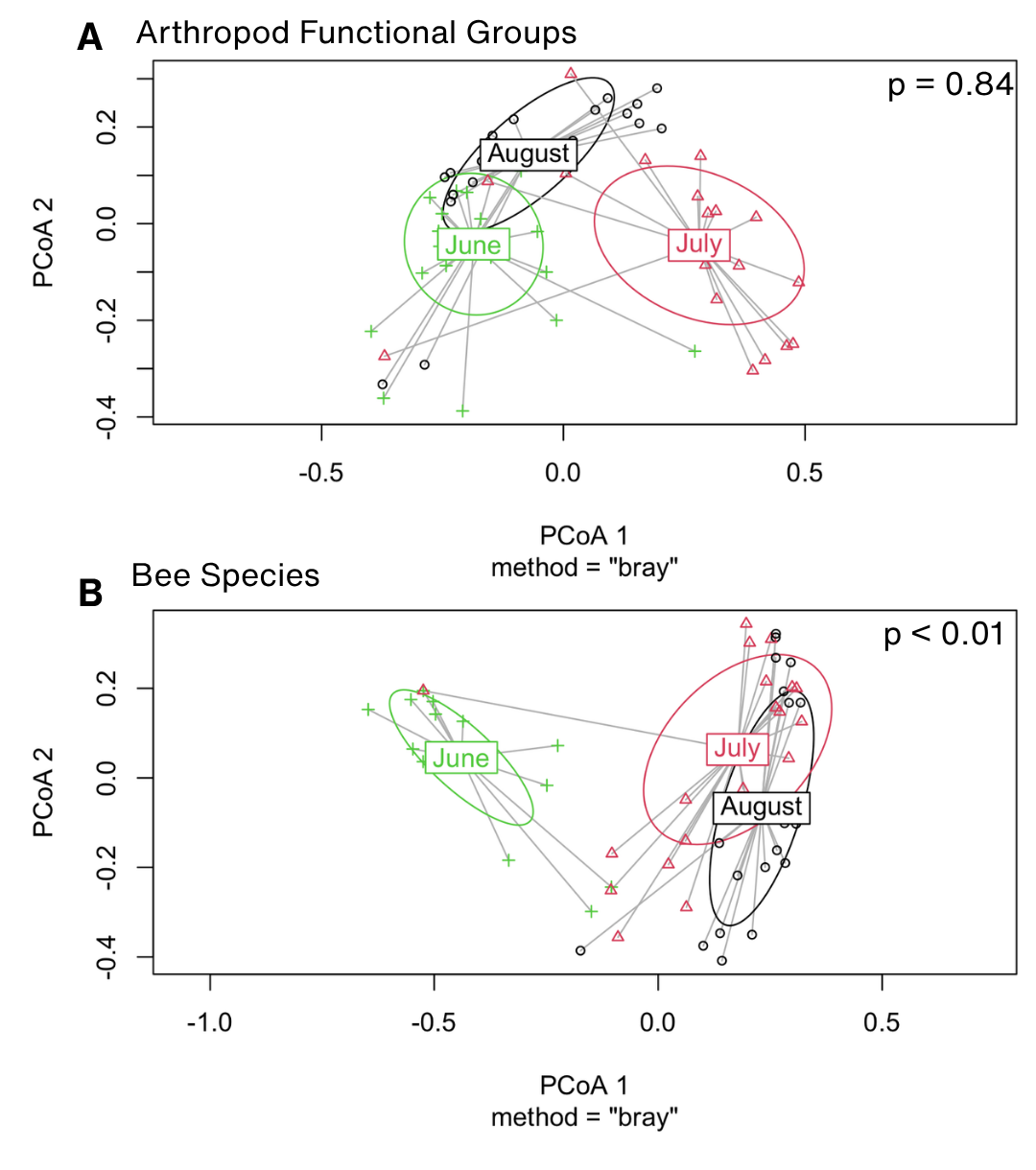

**Figure S1.3 PCoA plots of community composition.** Principal coordinates analysis (PCoA) of community composition based on Bray–Curtis dissimilarities for **(a)** arthropod functional groups and (**b)** bee species communities across months (June, July, August). Points represent individual sampling sites, colored by month. Ellipses denote 95% confidence intervals around group centroids, and line segments connect samples to their respective group centroids. Results of tests for homogeneity of multivariate dispersion (PERMDISP) are shown in each panel. Dispersion did not differ among months for arthropod functional groups (p = 0.84) but differed significantly for bee species communities (p < 0.01).

**Table S1.6** **Posterior summaries for total arthropod abundance.** Bayesian posterior summaries for the negative binomial regression of total arthropod abundance as a function of *wealth index* and *impervious surface*. *Month* was additionally included as a fixed effect, and *site* as a random effect. For each parameter, the table reports the posterior mean, standard error, 95% credible intervals, exponentiated estimates (back-transformed from the log scale), and the corresponding percent change in total arthropod abundance. Bolded values indicate significant population-level effects where credible intervals do not overlap zero.

| **Parameter** | **Estimate** | **Est. Error** | **Lower 95% CI** | **Upper 95% CI** | **e^β^** | **% change in abundance** |
| --- | --- | --- | --- | --- | --- | --- |
| Intercept | **4.23** | 0.11 | 4.02 | 4.45 | 68.63 | - |
| Impervious surface | **-0.50** | 0.10 | -0.68 | -0.31 | 0.61 | **-39.0** per +1 SD |
| Wealth Index | 0.08 | 0.09 | -0.10 | 0.26 | 1.09 | 8.7 |
| Month (July) | **0.85** | 0.11 | 0.63 | 1.07 | 2.35 | **134.9** |
| Month (June) | -0.20 | 0.11 | -0.43 | 0.02 | 0.81 | -18.5 |
| Site Intercept (Random Effect) | 0.34 | 0.09 | 0.19 | 0.54 | 1.41 | 40.7 |

**Table S1.7 Posterior summaries for pollinator, herbivore, and natural enemy abundance.** Bayesian posterior summaries for the negative binomial regressions of pollinators, herbivores, and natural enemies as a function of *wealth index* and *impervious surface*. *Month* was additionally included as a fixed effect, and *site* as a random effect. For each parameter, the table reports the posterior mean, standard error, 95% credible intervals, exponentiated estimates (back transformed from the log scale), and the corresponding percent change in estimated abundance. Bolded values indicate significant population-level effects where credible intervals do not overlap zero.

| **Parameter** | **Estimate** | **Est. Error** | **Lower 95% CI** | **Upper 95% CI** | **e^β^** | **% change in abundance** |
| --- | --- | --- | --- | --- | --- | --- |
| Pollinator Abundance | | | | | |  |
| Intercept | **3.13** | 0.16 | 2.82 | 3.44 | 22.85 | - |
| Impervious surface | **-0.51** | 0.14 | -0.78 | -0.24 | 0.60 | **-39.9** per +1 SD |
| Wealth Index | 0.08 | 0.12 | -0.15 | 0.32 | 1.08 | 8.3 |
| Month (July) | **0.39** | 0.19 | 0.02 | 0.76 | 1.48 | **47.5** |
| Month (June) | -1.36 | 0.22 | -1.77 | -0.92 | 0.26 | -74.3 |
| Site Intercept (Random Effect) | 0.31 | 0.15 | 0.03 | 0.62 | 1.36 | 36.2 |
| Herbivore Abundance | | | | | | |
| Intercept | **2.39** | 0.20 | 2.00 | 2.80 | 10.94 | - |
| Impervious surface | **-0.45** | 0.16 | -0.75 | -0.13 | 0.64 | **-36.4** per +1 SD |
| Wealth Index | 0.07 | 0.15 | -0.22 | 0.36 | 1.08 | 7.6 |
| Month (July) | **1.79** | 0.21 | 1.37 | 2.19 | 6.00 | **500.2** |
| Month (June) | 0.31 | 0.21 | -0.12 | 0.71 | 1.36 | 36.4 |
| Site Intercept (Random Effect) | 0.54 | 0.20 | 0.10 | 0.93 | 1.72 | 71.5 |
| Natural Enemy Abundance | | | | | | |
| Intercept | **2.87** | 0.12 | 2.63 | 3.12 | 17.56 | - |
| Impervious surface | **-0.41** | 0.09 | -0.60 | -0.23 | 0.66 | **-33.9** per +1 SD |
| Wealth Index | 0.08 | 0.08 | -0.08 | 0.24 | 1.08 | 8.2 |
| Month (July) | **0.52** | 0.16 | 0.20 | 0.83 | 1.68 | **68.1** |
| Month (June) | -0.21 | 0.16 | -0.52 | 0.10 | 0.81 | -19.0 |
| Site Intercept (Random Effect) | 0.18 | 0.11 | 0.01 | 0.41 | 1.19 | 19.4 |

**Table S1.8** **Posterior summaries for bee richness.** Bayesian posterior summaries for mixed-effects regressions of bee species richness, bee Shannon diversity, and bee Simpson diversity, as a function of *wealth index* and *impervious surface*. A smoothing spline function was added to wealth index to account for a non-linear relationship with the diversity outcomes. *Month* was additionally included as a fixed effect, and *site* as a random effect. Bolded values indicate significant population-level effects where credible intervals do not overlap zero. Note: the standard deviation of the smoothing spline (sds Wealth Index) is a hyperparameter, where credible intervals that do not overlap zero (bolded below) indicate significant non-linearity in the relationship.

| **Parameter** | **Estimate** | **Est. Error** | **Lower 95% CI** | **Upper 95% CI** |
| --- | --- | --- | --- | --- |
| Bee Species Richness | | | | |
| Intercept | **1.75** | 0.09 | 1.57 | 1.93 |
| Impervious Surface | **-0.34** | 0.09 | -0.51 | -0.17 |
| sWealth Index | -0.08 | 0.19 | -0.48 | 0.30 |
| Month (July) | **0.26** | 0.11 | 0.05 | 0.46 |
| Month (June) | **-0.39** | 0.12 | -0.64 | -0.15 |
| sds Wealth Index (Spline Hyperparameter) | **0.20** | 0.15 | 0.02 | 0.60 |
| Site Intercept (Random Effect) | 0.11 | 0.08 | 0.00 | 0.29 |
| Bee Shannon Diversity | | | | |
| Intercept | **1.17** | 0.09 | 0.98 | 1.35 |
| Impervious Surface | **-0.25** | 0.07 | -0.40 | -0.11 |
| sWealth Index | -0.05 | 0.18 | -0.40 | 0.32 |
| Month (July) | **0.30** | 0.11 | 0.07 | 0.51 |
| Month (June) | -0.06 | 0.11 | -0.29 | 0.17 |
| sds Wealth Index  (Spline Hyperparameter) | **0.16** | 0.14 | 0.01 | 0.53 |
| Site Intercept  (Random Effect) | 0.15 | 0.09 | 0.01 | 0.34 |
| σ (sigma) | 0.40 | 0.04 | 0.32 | 0.49 |
| Bee Simpson Diversity | | | | |
| Intercept | **0.93** | 0.09 | 0.75 | 1.11 |
| Impervious Surface | **-0.20** | 0.07 | -0.34 | -0.05 |
| sWealth Index | -0.03 | 0.17 | -0.36 | 0.33 |
| Month (July) | **0.28** | 0.11 | 0.06 | 0.48 |
| Month (June) | 0.10 | 0.11 | -0.13 | 0.31 |
| sds Wealth Index (Spline Hyperparameter) | 0.13 | 0.12 | 0.00 | 0.46 |
| Site Intercept (Random Effect) | 0.17 | 0.09 | 0.02 | 0.35 |
| σ (sigma) | 0.37 | 0.04 | 0.30 | 0.46 |

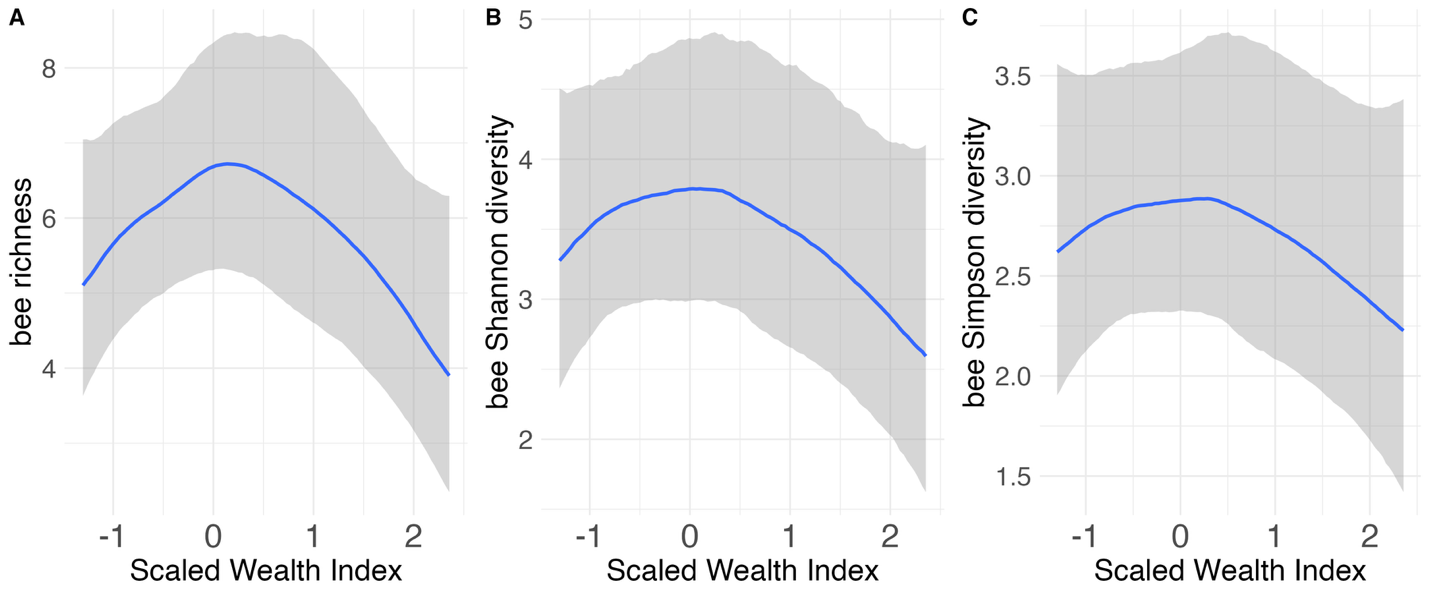

**Figure S1.4 Conditional effects of Wealth Index on bee biodiversity metrics.** Plots of the modeled relationships derived from Bayesian generalized additive models adjusting for the impervious surface cover and month. Panels show responses for **a)** bee species richness, **b)** bee Shannon diversity, and **c)** bee Simpson diversity, using a smoothing spline on Wealth Index to model non-linearities. The Wealth Index exhibited a hump-shaped, non-linear relationship with both bee richness (spline SD = 0.20, 95% CI [0.02,0.60]) and Shannon diversity (spline SD = 0.16, 95% CI [0.01,0.49]) while Simpson diversity showed a hump-shaped trend, though the credible interval for its smoothing parameter overlapped zero, suggesting an uncertain non-linear trend (spline SD = 0.13, 95% CI [0.00,0.46]). Shaded areas represent 95% credible intervals.

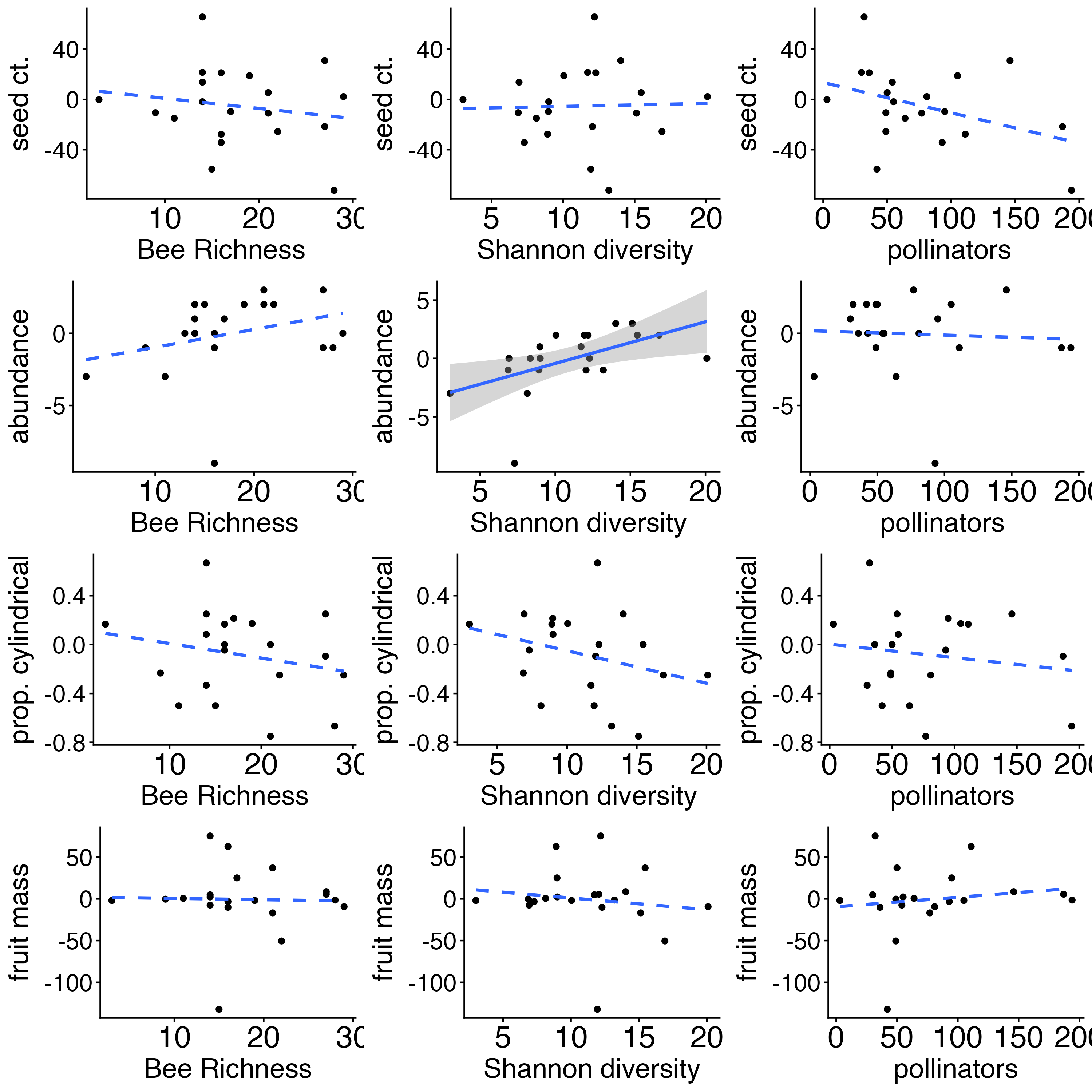

**Figure S1.5 Cucumber outcomes.** Scatterplots showing the difference between open-pollinated and hand-pollinated cucumber metrics (seed count, fruit abundance, proportion cylindrical, and fruit mass shown in top to bottom rows) averaged at each site as a function of *bee richness*, *bee Shannon diversity*, and *pollinator abundance* (left to right columns). Dashed lines indicate insignificant relationships, and solid lines indicate significant relationships.

**Table S1.9** **Posterior summaries for cucumber outcomes.** Cucumber outcomes as a function of arthropod predictors. All arthropod predictors were standardized/scaled prior to analysis. Values shown are posterior mean β estimates with 95% credible intervals in parentheses. Bolded values indicate significant effects where credible intervals do not overlap zero.

| Cucumber outcomes and arthropod predictors | | | | | | |
| --- | --- | --- | --- | --- | --- | --- |
| **Predictor** | **Δ Abundance (*Student t*)** | | **Δ Seeds**  **(*Student t*)** | | **Δ Mass**  **(*Student t*)** | **Δ Cylindrical (*Gaussian*)** |
| Intercept | 0.238 (-0.78, 1.15) | | -5.156 (-18.86, 8.08) | | 0.440 (-16.62, 17.43) | -0.078 (-0.23, 0.07) |
| Bee Shannon diversity (scaled) | **1.214 (0.20, 2.23)** | | 1.777 (-9.91, 12.97) | | -5.053 (-20.96, 11.13) | -0.095 (-0.24, 0.06) |
| Pollinator abundance (scaled) | -0.563 (-1.54, 0.47) | | -9.334 (-21.48, 3.78) | | 3.419 (-13.55, 20.79) | -0.079 (-0.24, 0.08) |
| Herbivore abundance (scaled) | 0.469 (-0.75, 1.72) | | 3.444 (-9.53, 16.73) | | 7.163 (-12.87, 26.50) | 0.124 (-0.06, 0.30) |
| Natural enemy abundance (scaled) | -0.016 (-1.25, 1.11) | | -0.923 (-13.32, 11.52) | | -7.401 (-27.78, 9.92) | -0.011 (-0.18, 0.16) |
| Additional model parameters | | | | | | |
| **Parameter** | | **Δ Abundance** | | **Δ Seeds** | **Δ Mass** | **Δ Cylindrical** |
| σ (sigma) | | 1.887 (0.95, 3.06) | | 28.666 (18.20, 42.53) | 33.417 (13.07, 54.31) | 0.351 (0.25, 0.51) |
| R^2^ | | 0.288 | | 0.161 | 0.131 | 0.247 |

**Table S1.10 Posterior summaries for structural equation model.** Bayesian posterior summaries for a two-part structural equation model for bee Shannon diversity and 𝚫 Cucumber abundance (insect-pollinated − hand-pollinated cucumbers). Bolded values indicate significant effects where credible intervals do not overlap zero.

| **Response Variable** | **Predictor** | **Estimate** | **SE** | **Lower 95% CI** | **Upper 95% CI** |
| --- | --- | --- | --- | --- | --- |
| **Bee Shannon diversity** | Intercept | -0.001 | 0.173 | -0.349 | 0.337 |
|  | Impervious surface (scaled) | **-0.526** | 0.188 | -0.890 | -0.150 |
|  | sWealth Index (scaled) | -0.190 | 0.380 | -0.930 | 0.600 |
|  | sds Wealth Index  (Spline Hyperparameter) | **0.260** | 0.200 | 0.010 | 0.760 |
|  | σ (sigma) | 0.843 | 0.144 | 0.613 | 1.173 |
| **𝚫 Cucumber abundance** | Intercept | 0.254 | 0.472 | -0.720 | 1.123 |
|  | Bee Shannon diversity (scaled) | **1.184** | 0.465 | 0.265 | 2.090 |
|  | σ (sigma) | 1.862 | 0.474 | 1.066 | 2.907 |

### **Appendix S2: Model Specifications and Bayesian Diagnostics**

### **Supplementary Methods: Bayesian model specification and priors**

Weakly informative priors for each model were set based on the data's spread and central tendency, and the likelihood link functions. Continuous predictors (wealth index and impervious surface) were centered and standardized (mean = 0, SD = 1) so that all regression slopes (β) were on a shared scale.

- **Intercept (α):** Represents the expected response value at average predictor values
- **Effect sizes (β):** Normal priors centered at zero with standard deviations reflecting expected effect scales. These priors act as weak conservative regularizers, penalizing excessively large slopes and reducing overestimation risk without imposing strict directionality.
- **Hurdle intercept** **(α_intercept_):** Specifies the baseline log-odds of observing a non-zero value at average predictor levels.
- **Group-level variation (SD_group_):** Group-level random effect variance. A larger SD value indicates more heterogeneity across groups.
- **Negative Binomial & Gamma shape (θ):** Accounts for overdispersion. A smaller value indicates a larger variance relative to the mean.
- **Residual error (𝜎):** Standard deviation of the residual error. Variation in the response variable unexplained by the predictors. A larger value indicates greater unexplained variance.
- **Degrees of freedom (𝛎):** The degrees of freedom parameter in a Student's t distribution, which determines how robust the model is to outliers. A small value indicates a higher weight is allowed to outliers; a large value indicates response values approaching a normal distribution.

**Table S2.1 Bayesian local and landscape response model priors.** All models demonstrated good convergence: R̂ < 1.01; Bulk ESS > 1000; Tail ESS > 1000 for all parameters.

| **Response Variable** | **Parameter** | **Prior** |
| --- | --- | --- |
| **Local Variables** | | |
| Tree and shrub cover (m²)  *hurdle gamma* | α | Normal (4, 1.5) |
|  | α_intercept_ | Normal (0, 1.5) |
|  | β | Normal (0, 1) |
|  | θ | Gamma (2, 0.5) |
| Garden flora richness  *gaussian* | α | Normal (60, 15) |
|  | β | Normal (0, 5) |
|  | 𝜎 | Exponential (0.05) |
| Garden flora Shannon diversity  *gaussian* | α | Normal (45, 15) |
|  | β | Normal (0, 5) |
|  | 𝜎 | Exponential (0.08) |
| Bare ground (m²)  *hurdle gamma* | α | Normal ( 2.5, 1.5) |
|  | α_intercept_ | Normal (0, 1.5) |
|  | β | Normal (0, 1) |
|  | θ | Gamma (2, 0.5) |
| Flora density (species/m²)  *lognormal* | α | Normal (-1, 1) |
|  | β | Normal (0, 1) |
|  | 𝜎 | Exponential (2) |
| **Landscape Variables** | | |
| Tree canopy cover (500 m)  *gaussian* | α | Normal (12, 5) |
|  | β | Normal (0, 5) |
|  | 𝜎 | Exponential (0.1) |
| Landscape flora rarefied richness (1000 m)  *gaussian* | α | Normal (115, 30) |
|  | β | Normal (0, 20) |
|  | 𝜎 | Exponential (0.03) |
| Landscape flora rarefied Shannon diversity (1000 m)  *gaussian* | α | Normal (80, 25) |
|  | β | Normal (0, 15) |
|  | 𝜎 | Exponential (0.04) |
| Water & wetland (1000 m)  *hurdle gamma* | α | Normal (-3.2, 0.5) |
|  | α_intercept_ | Normal (0, 1.5) |
|  | β | Normal (0, 1) |
|  | θ | Gamma (10, 2) |
| Shrub & grassland (1000 m)  *hurdle gamma* | α | Normal (-3.9, 0.5) |
|  | α_intercept_ | Normal (0, 1.5) |
|  | β | Normal (0, 1) |
|  | θ | Gamma (10, 2) |

**Table S2.2 Bayesian arthropod abundance and bee diversity model priors.** All models demonstrated good convergence: R̂ < 1.01; Bulk ESS > 1000; Tail ESS > 1000 for all parameters.

| **Response Variable** | **Parameter** | **Prior** |
| --- | --- | --- |
| Total arthropod abundance  *negative binomial* | α | Normal (log80, 0.5) |
|  | β | Normal (0, 1) |
|  | SD_group_ | Exponential (1) |
|  | θ | Gamma (2, 0.1) |
| Pollinator abundance  *negative binomial* | α | Normal (log10, 0.5) |
|  | β | Normal (0, 0.5) |
|  | SD_group_ | Exponential (1) |
|  | θ | Gamma (2, 0.1) |
| Herbivore abundance  *negative binomial* | α | Normal (log20, 0.5) |
|  | β | Normal (0, 0.5) |
|  | SD_group_ | Exponential (1) |
|  | θ | Gamma (2, 0.1) |
| Natural enemy abundance  *negative binomial* | α | Normal (log18, 0.4) |
|  | β | Normal (0, 0.4) |
|  | SD_group_ | Exponential (1) |
|  | θ | Gamma (2, 0.1) |
| Bee species richness  *poisson* | α | Normal (1.38, 0.5) |
|  | β | Normal (0, 0.25) |
|  | SD_spline_ | Normal (0, 0.5) |
|  | SD_group_ | Exponential (2) |
| Bee Shannon Diversity  *lognormal* | α | Normal (1.25, 0.5) |
|  | β | Normal (0, 0.25) |
|  | 𝜎 | Exponential (1) |
|  | SD_spline_ | Normal (0, 0.5) |
|  | SD_group_ | Exponential (2) |
| Bee Simpson Diversity  *lognormal* | α | Normal (1.10, 0.5) |
|  | β | Normal (0, 0.25) |
|  | 𝜎 | Exponential (1) |
|  | SD_spline_ | Normal (0, 0.5) |
|  | SD_group_ | Exponential (2) |

**Table S2.3 Cucumber outcomes model priors.** All models demonstrated good convergence: R̂ < 1.01; Bulk ESS > 1000; Tail ESS > 1000 for all parameters.

| **Response** | **Parameter** | **Prior** |
| --- | --- | --- |
| Δ Cucumber abundance  *student t* | α | Normal (0, 5) |
|  | β | Normal (0, 2) |
|  | 𝜎 | Exponential (0.2) |
|  | 𝜈 | Gamma (2, 0.1) |
| Δ No. of Seeds  *student t* | α | Normal (0, 40) |
|  | β | Normal (0, 10) |
|  | 𝜎 | Exponential (0.03) |
|  | 𝜈 | Gamma (2, 0.1) |
| Δ Cucumber fruit mass  *student t* | α | Normal (0, 75) |
|  | β | Normal (0, 20) |
|  | 𝜎 | Exponential (0.02) |
|  | 𝜈 | Gamma (2, 0.1) |
| Δ Proportion cylindrical  *gaussian* | α | Normal (0, 0.5) |
|  | β | Normal (0, 0.2) |
|  | 𝜎 | Exponential (4) |

**Table S2.4 Bayesian SEM model priors.** All models demonstrated good convergence: R̂ < 1.01; Bulk ESS > 1000; Tail ESS > 1000 for all parameters.

| **Response Variable** | **Parameter** | **Prior** |
| --- | --- | --- |
| Bee Shannon diversity  *gaussian* | α | Normal (0, 0.5) |
|  | β | Normal (0, 0.5) |
|  | SD_group_ | Normal (0, 0.5) |
|  | 𝜎 | Exponential (2) |
| 𝚫 Cucumber abundance  *student t* | α | Normal (0, 5) |
|  | β | Normal (0, 2) |
|  | 𝜎 | Exponential (0.2) |
|  | 𝜈 | Gamma (2, 0.1) |

**Posterior Predictive Plots**

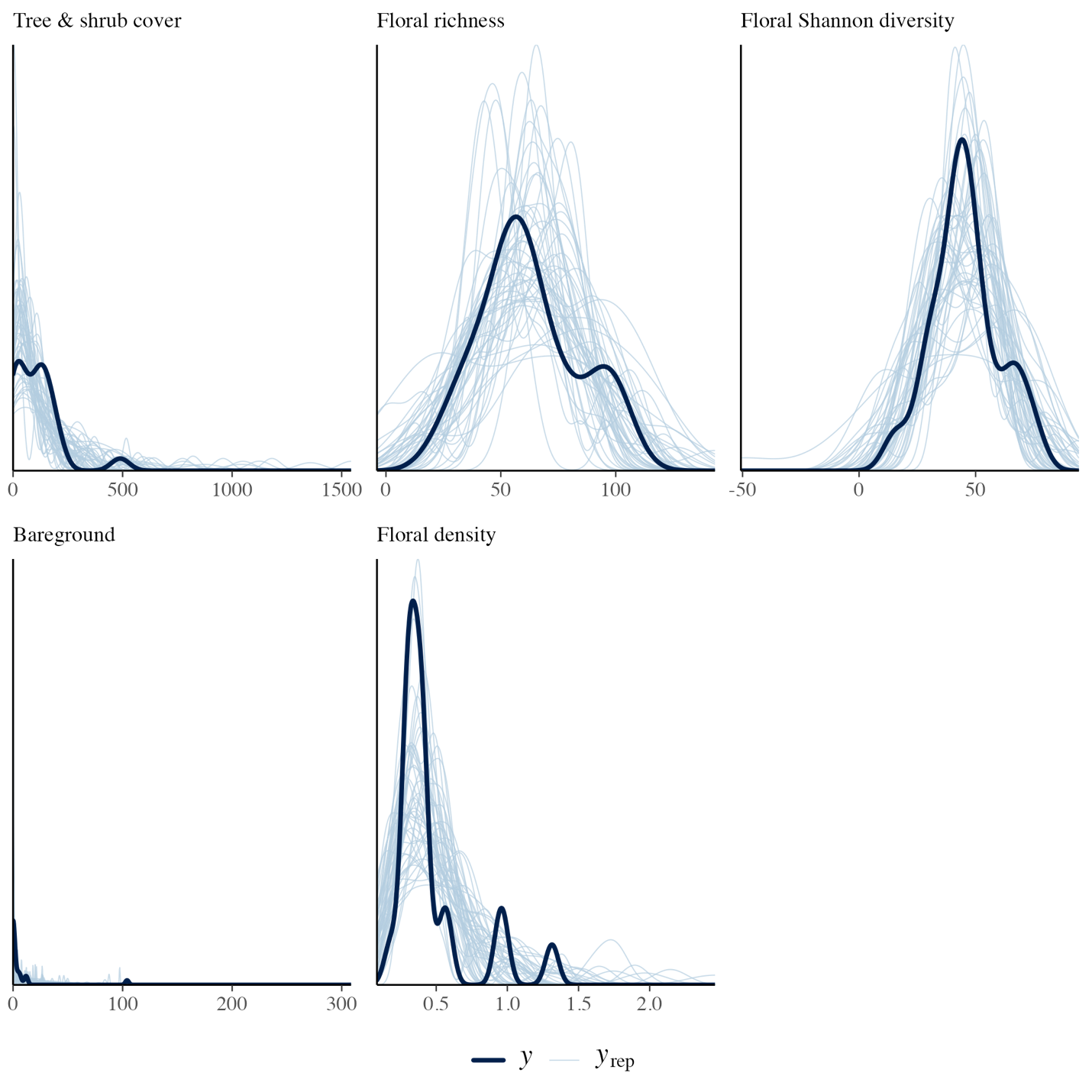

**Figure S2.1 Posterior predictive plots of local habitat quality outcome variables.** Posterior outcomes were simulated 50 times using the parameter estimates from each model.

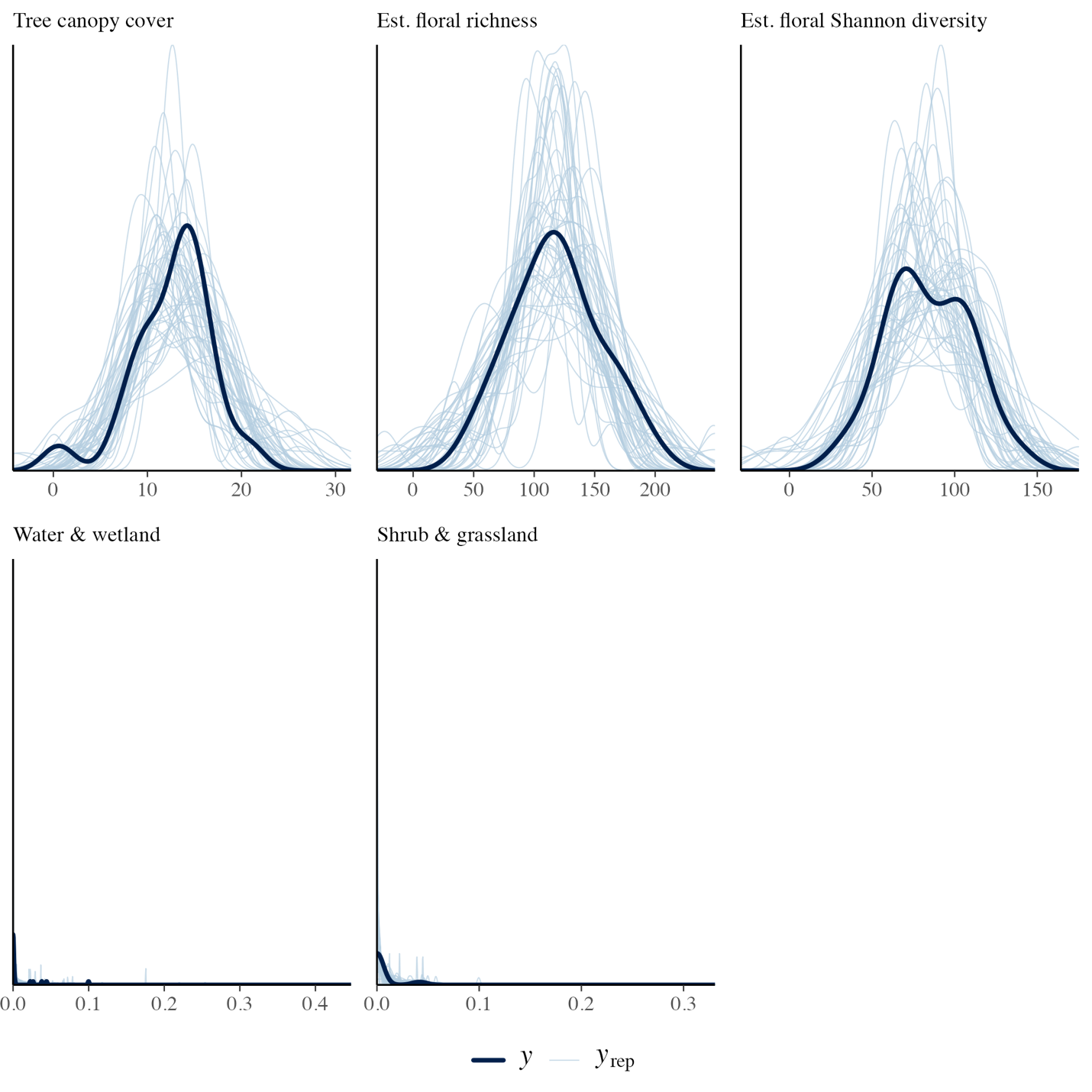

**Figure S2.2 Posterior predictive plots of landscape habitat quality outcome variables.** Posterior outcomes were simulated 50 times using the parameter estimates from each model.

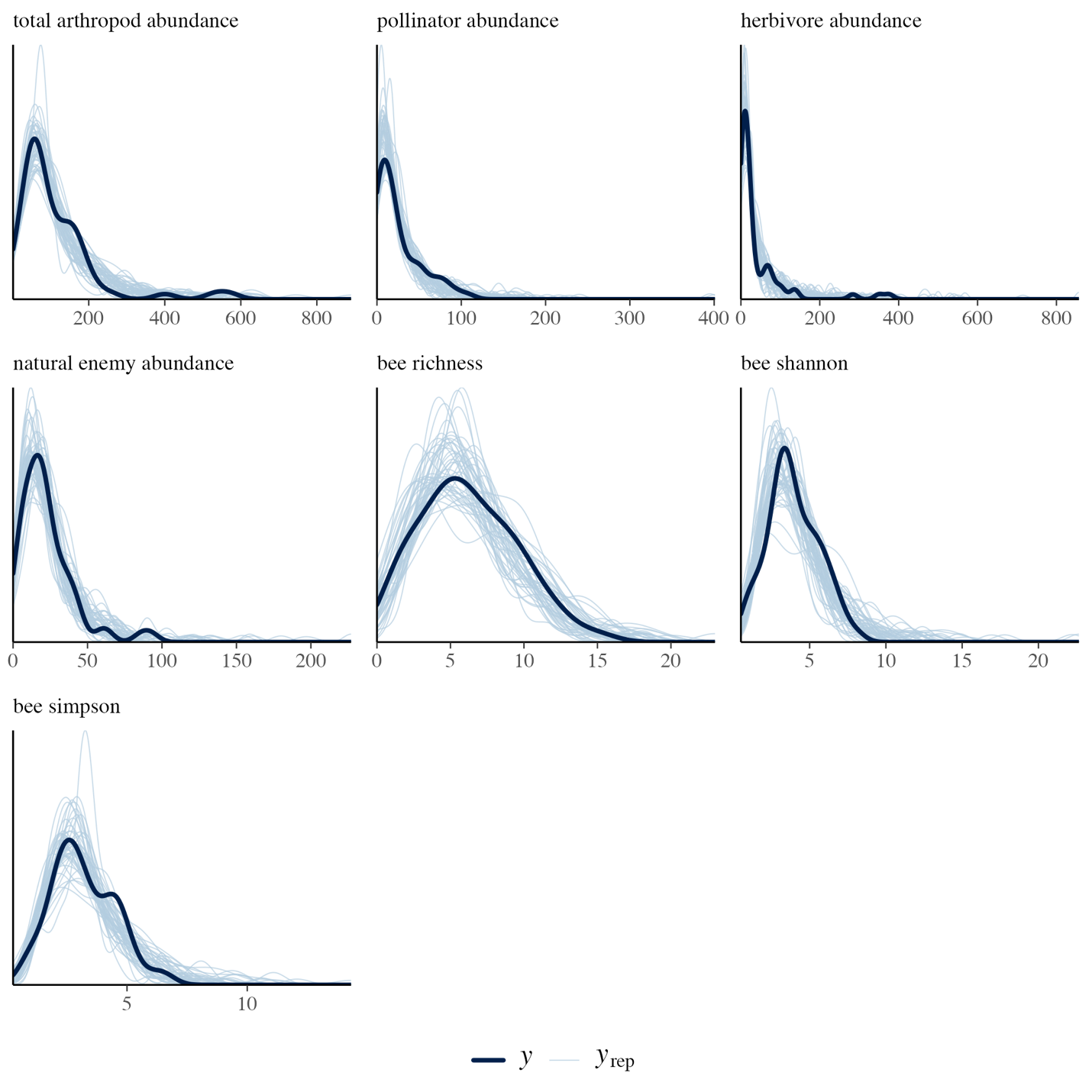

**Figure S2.3 Posterior predictive plots of arthropod outcome variables.** Posterior outcomes were simulated 50 times using the parameter estimates from each model.

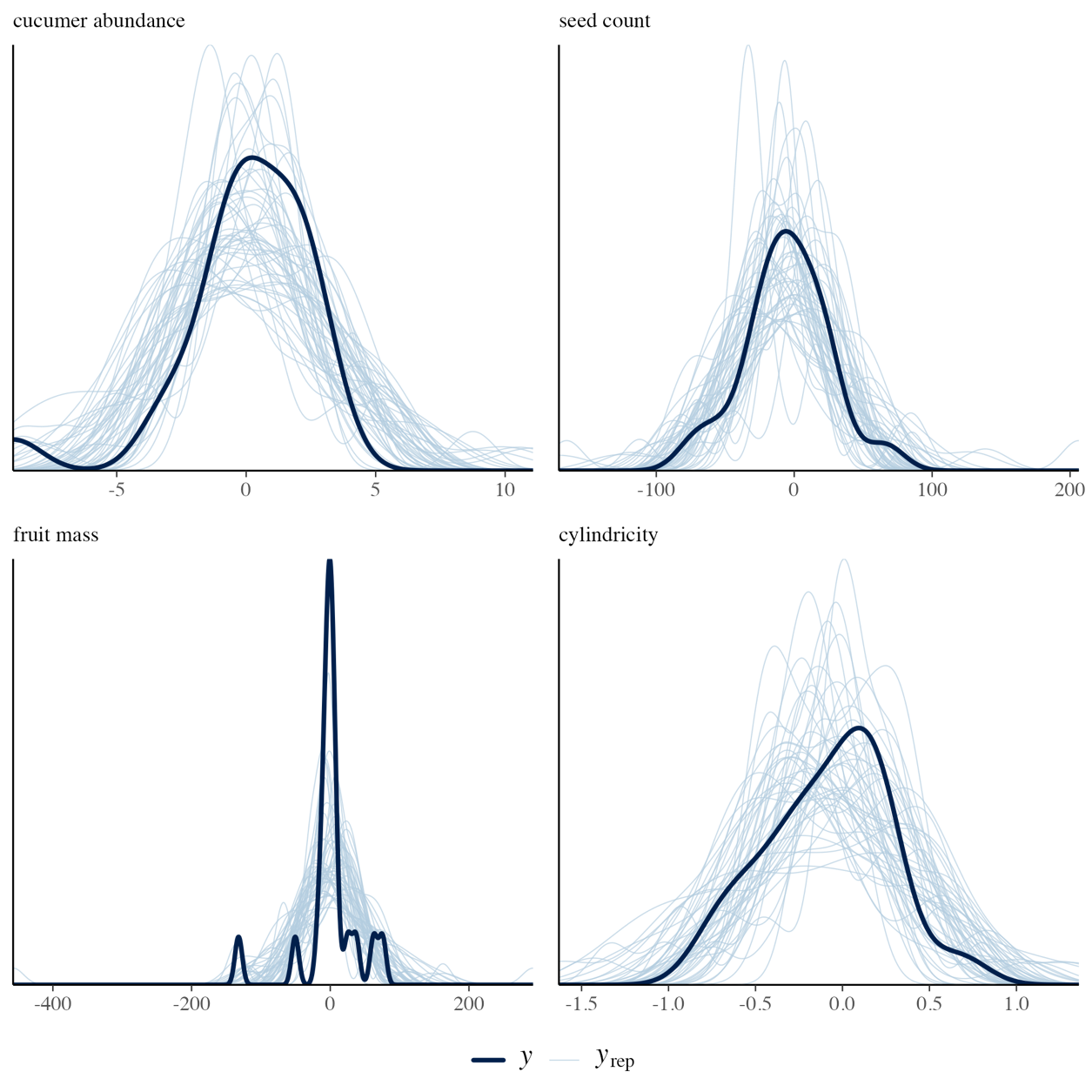

**Figure S2.4 Posterior predictive plots of cucumber outcome variables.** Posterior outcomes were simulated 50 times using the parameter estimates from each model.

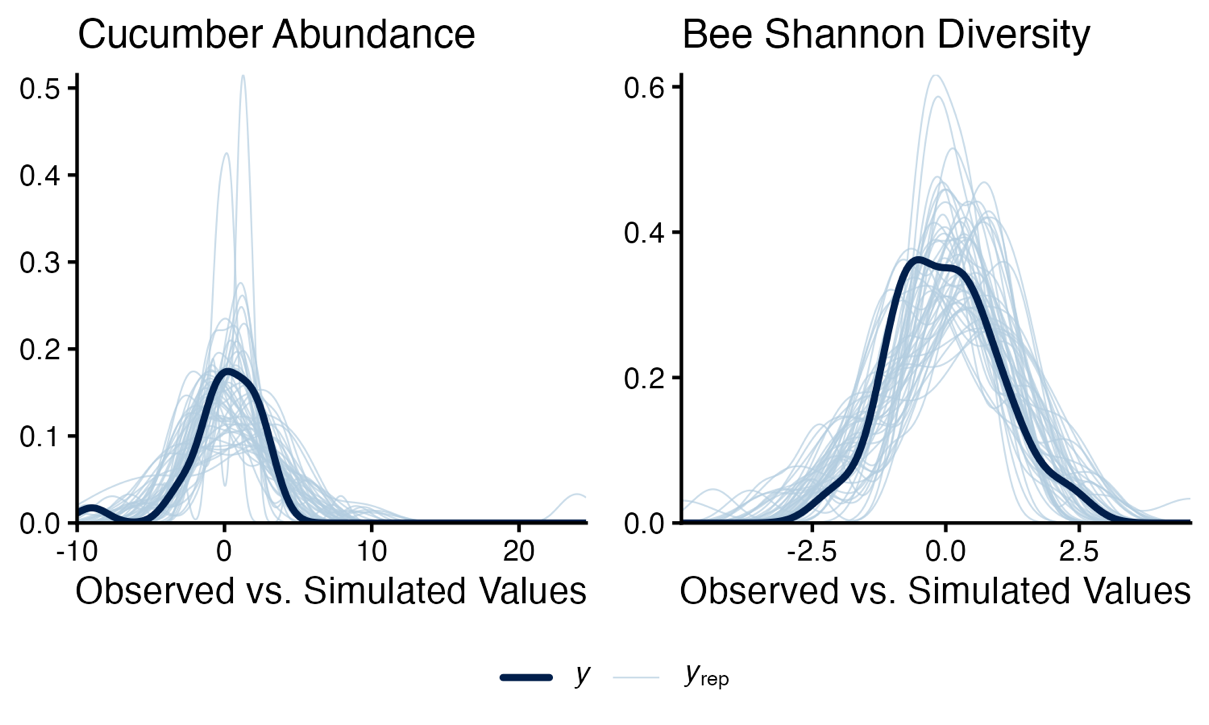

**Figure S2.5 Posterior predictive plots of structural equation model outcomes.** Posterior outcomes were simulated 50 times using the parameter estimates from each model.
